# Multi-scale hierarchical neural network models that bridge from single neurons in the primate primary visual cortex to object recognition behavior

**DOI:** 10.1101/2021.03.01.433495

**Authors:** Tiago Marques, Martin Schrimpf, James J. DiCarlo

## Abstract

Primate visual object recognition relies on the representations in cortical areas at the top of the ventral stream that are computed by a complex, hierarchical network of neural populations. While recent work has created reasonably accurate image-computable hierarchical neural network models of those neural stages, those models do not yet bridge between the properties of individual neurons and the overall emergent behavior of the ventral stream. One reason we cannot yet do this is that individual artificial neurons in multi-stage models have not been shown to be functionally similar to individual biological neurons. Here, we took an important first step by building and evaluating hundreds of hierarchical neural network models in how well their artificial single neurons approximate macaque primary visual cortical (V1) neurons. We found that single neurons in certain models are surprisingly similar to their biological counterparts and that the distributions of single neuron properties, such as those related to orientation and spatial frequency tuning, approximately match those in macaque V1. Critically, we observed that hierarchical models with V1 stages that better match macaque V1 at the single neuron level are also more aligned with human object recognition behavior. Finally, we show that an optimized classical neuroscientific model of V1 is more functionally similar to primate V1 than all of the tested multi-stage models, suggesting room for further model improvements with tangible payoffs in closer alignment to human behavior. These results provide the first multi-stage, multi-scale models that allow our field to ask precisely how the specific properties of individual V1 neurons relate to recognition behavior.

**Highlights:**

- Image-computable hierarchical neural network models can be naturally extended to create hierarchical “brain models” that allow direct comparison with biological neural networks at multiple scales – from single neurons, to population of neurons, to behavior.
- Single neurons in some of these hierarchical brain models are functionally similar to single neurons in macaque primate visual cortex (V1)
- Some hierarchical brain models have processing stages in which the entire distribution of artificial neuron properties closely matches the biological distributions of those same properties in macaque V1
- Hierarchical brain models whose V1 processing stages better match the macaque V1 stage also tend to be more aligned with human object recognition behavior at their output stage

## Introduction

The primate ventral visual stream, a complex network of hierarchically-organized cortical areas, has been shown to support visually-guided behaviors (Felleman & Van Essen, 1991; Mishkin, Ungerleider, & Macko, 1983). One such particularly important behavior is core object recognition -- i.e., the ability to rapidly (~200ms) identify objects in the central visual field (DiCarlo, Zoccolan, & Rust, 2012; Fabre-Thorpe, Richard, & Thorpe, 1998). Understanding the computations and neuronal mechanisms underlying this challenging visual behavior has been a major goal in systems neuroscience (DiCarlo & Cox, 2007). A critical step towards this goal is the development of accurate, multi-stage, multi-scale models that can bridge between the properties of individual cells and phenomena at each of the ventral stream processing stages to the supported visually-guided behaviors, such as core object recognition. These multi-stage, multi-scale models would, for example, allow us to begin to understand how functional properties at the cellular level, where we can most precisely measure and manipulate the system, give rise to visually intelligent behavior. Successful multi-scale models must be simultaneously accurate at both the individual neuron level, at the neuronal population level, and at the behavioral level. The work presented here is one step toward that greater goal.

Prior work has shown that hierarchical networks of artificial neural populations can, when properly connected, quite closely approximate object recognition behavioral patterns that are driven by the ventral visual stream, a similarly organized deep hierarchy of biological neurons (Rajalingham et al., 2018; Schrimpf et al., 2018). In addition, this same model family has achieved unparalleled success in explaining the response patterns of individual neurons along the ventral stream areas (Bashivan, Kar, & DiCarlo, 2019; Cadena et al., 2019; Kar, Kubilius, Schmidt, Issa, & DiCarlo, 2019; Schrimpf et al., 2018; Yamins et al., 2014). Thus, we and others have proposed that these models may serve as multi-stage, multi-scale models of the mechanisms of object recognition – causally bridging from single neuron responses at multiple stages of the ventral stream to the observed recognition behavioral patterns (Kriegeskorte, 2015; Richards et al., 2019; Schrimpf et al., 2020; Yamins & DiCarlo, 2016). However, this model to brain congruency has not been without criticism. In particular, these models often contain critical unspecified parameters such as: the mapping between their input and a physical field-of-view, and the commitment of specific model stages to specific brain regions. In addition, when using these models to predict neuronal responses, researchers (including some of the current authors) have relied on fitting methods that linearly combine thousands of features, or model neurons, to explain the responses of individual biological neurons (Saxe, Nelli, & Summerfield, 2021; Serre, 2019). The lack of a pre-specified model-to-brain mapping means that hierarchical neural networks cannot be yet considered to be accurate multiscale models of the neural mechanisms of object recognition. For example, the congruency tests typically do not require that the individual artificial single neurons are aligned with individual biological neurons within the proposed congruent cortical area(s).

To address this limitation, we here hypothesized that these hierarchical models of artificial neurons might be modified to become accurate, multi-scale models of the neural mechanisms of visual object recognition. To investigate this, we first created a range of new candidate “brain models” by using existing base model architectures and exploring two key model parameters that are biologically critical: model field-of-view and model processing stage proposed to correspond to primate V1. We then explicitly mapped single artificial neurons in each of these hierarchical models to single biological neurons in the primate primary visual cortex (area V1) in a one-to-one mapping commitment. Specifically, we avoided the usual model-to-brain fitting procedure and we instead sought to test the hypothesis that single neurons in a candidate model layer (i.e. a specific processing stage of a candidate hierarchical model) correspond to single neurons in the macaque V1, and that the entire artificial neural population at that same model layer corresponds to the entire V1 neuronal population. That is, we asked if any models were explicitly well matched to primate V1 at both the single neuron level and the population level. We were encouraged to pursue this approach to modeling V1 in part because of prior work demonstrating that one such hierarchical network model contains neural representations which, when linearly combined using a regression approach, can reasonably accurately predict the response patterns of V1 (Cadena et al., 2019).

To compare models with primate V1 in this way, we performed in-silico neurophysiological experiments in hundreds of these V1 candidate brain models to measure 22 single neuron response properties that have been previously quantified, such as those related to orientation and spatial frequency tuning and surround and texture modulation, and compared their distributions to those in macaque V1 from available published studies (Cavanaugh, Bair, & Movshon, 2002; De Valois, Albrecht, & Thorell, 1982; De Valois, Yund, & Hepler, 1982; Freeman, Ziemba, Heeger, Simoncelli, & Movshon, 2013; Ringach, Shapley, & Hawken, 2002; P. H. Schiller, Finlay, & Volman, 1976; Ziemba, Freeman, Movshon, & Simoncelli, 2016).

We found that randomly-sampled single artificial neurons in the V1-layers of certain hierarchical brain models have response characteristics that are surprisingly similar to those of single neurons in macaque V1. We also found that the population distributions of response properties also very closely match the biological distributions of those same response properties. Since all of these hierarchical models were multi-stage candidate models of the entire ventral stream and its supported object recognition behavior, we then asked: Do ventral stream models that better align with biological V1 at their proposed V1 processing stage also better align with the behavioral patterns of human core object recognition? Indeed, we found that hierarchical models with a V1 stage that better matched macaque V1, had behavioral “output” that was more closely matched to human behavior. Thus, this work describes, for the first time, image-computable, multi-stage models of the primate visual ventral stream that bridge from single neurons in V1, the first visual cortical area, all the way to object recognition behavior.

Importantly however, we found that no evaluated ventral stream model was able to perfectly account for all the V1 response properties and all tested models underperformed when compared to an optimized classical neuroscientific model of V1. This shows that the causal, multi-scale models of the ventral stream developed here can be further improved, and argues that improvements -- even at just the V1 processing stage -- will lead to better causal models of human object recognition behavior.

## Results

Our overarching goal is to build accurate, multi-stage, multi-scale models of how the primate ventral visual stream supports object recognition behaviors. By definition, such models must be accurate at the level of single neurons and the level of populations of such neurons (multi-scale), as well as at all ventral stream stages and ventral stream behaviors (multi-stage). In this work, we focus on primate visual area V1 and we evaluate how well specific hierarchical artificial neural networks (ANNs), some of which are the current leading models of the ventral visual stream (see Schrimpf et al., 2018 and accompanying website for the current leading models; Cadena et al., 2019; Kubilius et al., 2019; Yamins et al., 2014), directly align at the level of V1 single neurons. Contrary to prior approaches that used fitting high dimensional feature spaces in the models to the responses of relatively small neuronal populations (Cadena et al., 2019) we here tested the even stronger hypothesis that single neurons in variants of the existing hierarchical ventral stream models may qualitatively and quantitatively align with single neuron functional properties in macaque V1 in a one-to-one manner (see Fig. 1, bottom).

**Figure 1.**
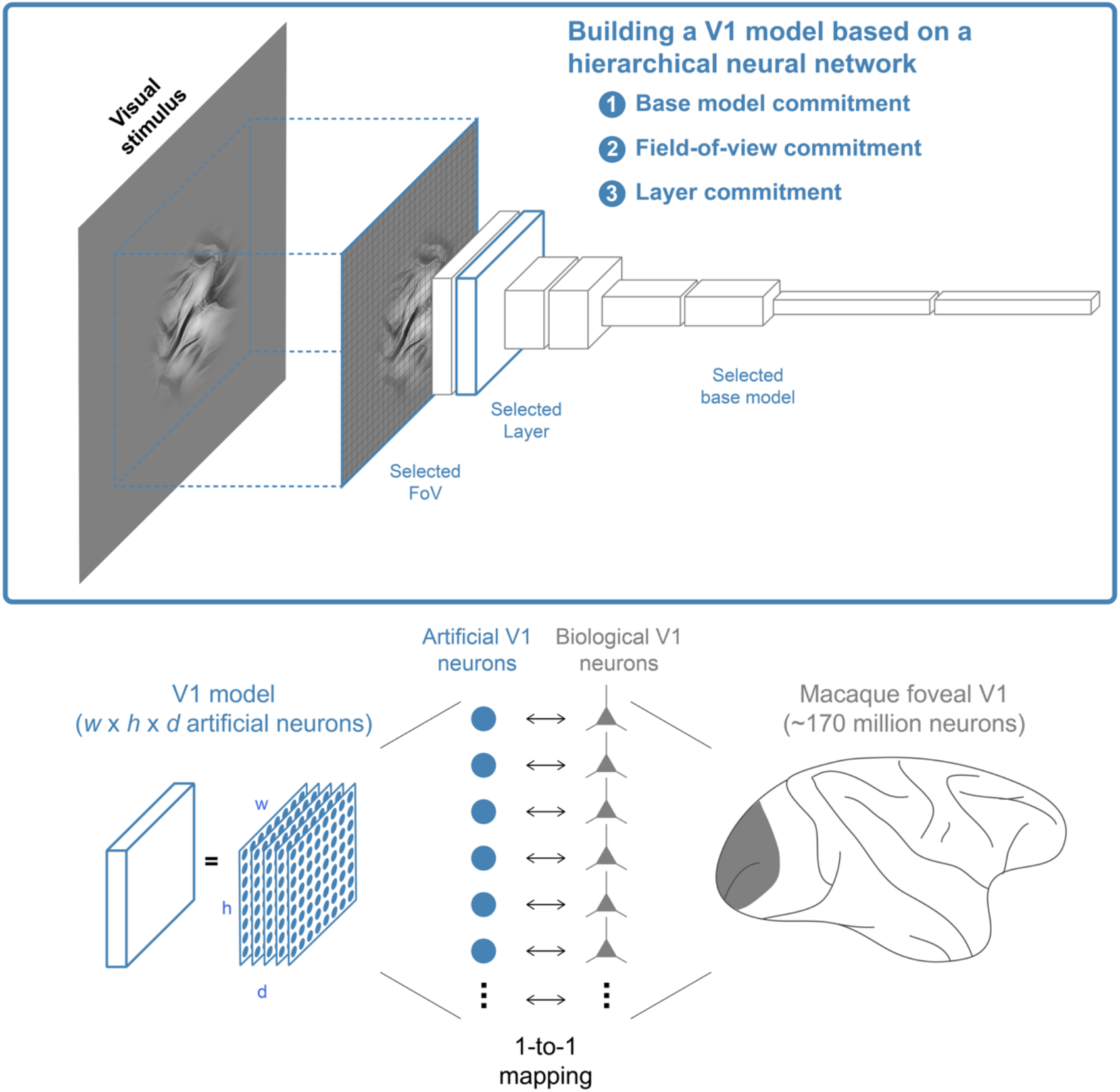
Building V1 models at the level of single neurons using hierarchical neural networks. Top, building a candidate model of macaque V1 involves three steps: (1) the choice of a base model defined by its architecture and synaptic weights, (2) the choice of the field-of-view (FoV) in physical units (degrees), and (3) the choice of the processing stage, i.e. layer, to map to V1. Bottom, the model of V1 based on a hierarchical neural network is a convolutional layer containing *w* × *h* × *d* neurons, where *w*, *h*, and *d*, are the width, height, and number of features, respectively. This modeling framework assumes a one-to-one mapping at the level of single neurons, i.e., each artificial neuron in the model corresponds to a putative biological neuron in macaque foveal V1.

To test this hypothesis, we developed a three-step approach for building hundreds of candidate models of V1 using specific, hierarchical ANNs that are already among the leading models of the ventral visual stream (Figure 1, top). First, we choose a base model (Schrimpf et al., 2020) consisting of a hierarchical network architecture and all its synaptic weights which are typically obtained by standard training on the object classification ImageNet dataset, though we also used models with their weights optimized differently (see Methods). Second, since the base model’s input is solely defined by its resolution in pixels (the model’s input sensors, 224×224 in all the models used) with no connection to physical quantities, we specified the region of visual space (in degrees) that corresponds to the model’s input and we termed that the field-of-view (FoV). In this study we considered multiple model FoVs. Relative to models with a smaller FoV, models with a larger FoV have the same number of input sensors, but each individual sensor integrates over a larger spatial extent, resulting in a larger combined sampled spatial extent (i.e. a larger FoV). Third, we assign all the artificial neurons within a specific layer of the hierarchical model as a candidate model of the macaque V1 neural population. Due to the convolutional architecture of the neural networks used, each model layer consists of multiple feature spatial maps and thus each candidate V1 contains *w* × *h* × *d* artificial neurons (range 10K-3M artificial neurons). To obtain a one-to-one mapping of artificial neurons to biological neurons, we discard information about each neuron’s spatial location and feature number and treat it as a putative single neuron in foveal macaque V1. In other words, for each candidate V1 we randomly sample artificial neurons from this pool as if we were randomly sampling individual neurons with a recording electrode. We then quantify the response properties of these individually sampled artificial individual neurons and compare them with analogous measurements of individual biological V1 neurons from multiple experiments.

In total, we considered: (1) 13 different base models including AlexNet (Krizhevsky, Sutskever, & Geoffrey E., 2012), VGG (Simonyan & Zisserman, 2015), ResNet (He, Zhang, Ren, & Sun, 2016), CORnet (Kubilius et al., 2019), and bagnet (Brendel & Bethge, 2019); (2) four different FoVs (between 4 and 10 degrees); and (3) multiple early and intermediate layers for each base model. This resulted in 736 candidate V1 models (see methods for a complete description).

### Single artificial neurons in some hierarchical networks have response patterns that are qualitatively similar to those of single neurons in macaque V1

Over the last several decades, responses of individual neurons in macaque V1 have been extensively characterized using different types of parametric stimuli such as gratings with varying phase, orientation, spatial frequency (SF) and size, and naturalistic textures and noise images (Figure 2B,C). Simple cells show responses strongly modulated by the phase of gratings while complex cells are invariant to this stimulus property (Skottun et al., 1991). Furthermore, V1 neurons vary widely in their orientation (De Valois, Yund, et al., 1982; Ringach et al., 2002; Peter H Schiller, Finlay, & Volman, 1976) and SF selectivities (De Valois, Albrecht, et al., 1982; P. H. Schiller et al., 1976) selectivities, receptive field (RF) sizes, and the degree to which stimuli outside their RFs modulate their responses (Cavanaugh et al., 2002; H. E. Jones, Grieve, Wang, & Sillito, 2001; Kapadia, Ito, Gilbert, & Westheimer, 1995; Lamme, 1995; Sceniak, Ringach, Hawken, & Shapley, 1999). Finally, V1 neurons tend to respond similarly to texture stimuli and noise images with matching spatially averaged orientation and SF structure (Freeman et al., 2013; Ziemba et al., 2016). In prior experimental work, these characteristics of neuronal responses were quantified by calculating response properties (Figure 2C) such as: the F1/F0 ratio (ratio of the first harmonic and the DC component of responses to drifting gratings, also known as phase modulation ratio), preferred orientation and circular variance (CV; quantifies how selective the responses to different orientations are), peak SF and SF bandwidth, grating summation field (GSF; size of the stimulus for which the response is maximized; related to the size of the excitatory component of the RF) and surround suppression index (SSI, quantifies how much responses are suppressed by stimuli outside the classical RF), and texture modulation index (TMI, quantifies how much stronger neurons respond to naturalistic textures versus noise images).

**Figure 2.**
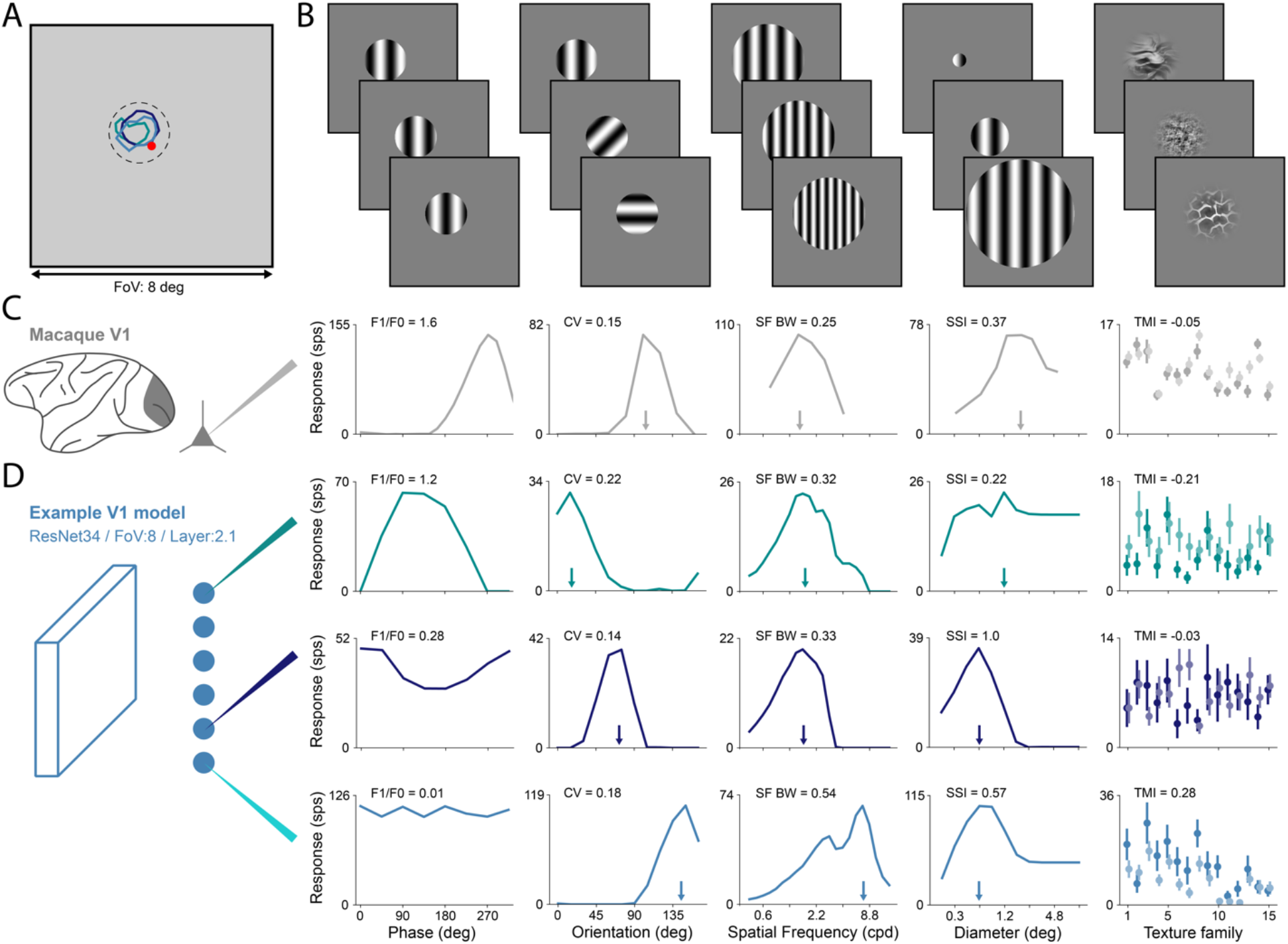
Single neurons in certain V1 models show responses similar to those of neurons in macaque V1. **A.** Field-of-view of example V1 model with 8 deg. Red circle shows center of gaze. Dashed circle represents the location of a stimulus with a circular aperture (2deg diameter). Colored contours show the receptive field locations of three example neurons aligned with the stimulus center. **B.** Example stimuli used in the in-silico neurophysiology characterization of single neurons in V1. From left to right: gratings with varying phase, gratings with varying orientation, gratings with varying spatial frequency, gratings with varying size, and naturalistic texture and noise images. **C.** Example responses of neurons in macaque V1. From left to right: phase, orientation, spatial frequency, and size tuning curves, and responses to naturalistic textures (dark tone) and spectrally matched noise images (light tone). Plots are vertically aligned with the corresponding example stimuli on B. Responses are taken from published studies and each plot corresponds to a different neuron. Values of example single neuron properties calculated from these responses are shown at the top of each corresponding plot (phase modulation ratio, circular variance, spatial-frequency bandwidth, surround suppression index, and texture modulation index). Arrows indicate the preferred orientation, peak spatial frequency, and grating summation field in their respective plots. **D.** Similar to C but for three example neurons from a neural network V1 model (Layer2.1 of ResNet34 with a FoV of 8deg). The plots on each row correspond to the same neuron with the receptive field shown in A with matching color. Within the same layer of the hierarchical neural network model, single neurons exhibit very different response characteristics. The neuron in the top row has a simple-cell like response with a strong phase modulation while the other two neurons show more complex-like responses. All neurons are strongly orientation and spatial-frequency selective but with different preferences and bandwidths. Neurons show different amounts of surround suppression and texture modulation.

Using the V1 candidate models previously described, we performed a series of in silico recordings to characterize the responses of their single neurons. After mapping the RFs of individual model neurons by presenting small gratings at different locations (Figure 2A; methods), we recorded their visual responses to the presentation of stimuli typically used to study macaque V1 (Figure 2B,D). We found that single neurons in some V1 candidate models show responses to visual stimuli that are similar to single neurons in macaque V1, allowing us to calculate response properties exactly the same way as with neurophysiology data (Figure 2D). Like in macaque V1, single artificial neurons within the same V1 model vary widely in their responses. For example, they vary in their selectivity to phase, orientation and SF of gratings, RF size, and in how their responses are inhibited by the presence of surrounding stimuli (Figure 2D).

In addition to the intra-model variability, we found that median single neuron response properties also vary considerably across alternative candidate V1 models. This intermodel variation is driven by the choice of: base model, field-of-view (FoV), and the layer (Supplementary Figure 1). Some response properties vary with these V1 model choices in an intuitive way. For example, a V1 model’s median neuronal RF size (as determined by the GSF) increases with the layer depth and with the FoV size: V1 candidate models selected from deep layers in the base model contain neurons that can potentially integrate input from larger portions of the visual field (relative to more shallow layers), and V1 candidate models with larger FoVs can potentially integrate over larger portions of visual space as measured in degrees (Supplementary Figure 1). Similarly, a V1 model’s median neuronal peak preferred spatial frequency decreases with increasing FoV. On the other hand, other response properties, which show a strong dependency on the layer depth, are not particularly affected by the FoV. For some base models, such as ResNet-34, circular variance increases monotonically with layer depth, which is analogous to a decrease in the number of orientation selective neurons observed along the primate ventral stream (Matteucci, Marotti, Riggi, Rosselli, & Zoccolan, 2019). Similarly, texture modulation index also increases with layer depth which once again is also observed along the primate ventral stream hierarchy (Freeman et al., 2013; Laskar, Giraldo, & Schwartz, 2018). An identical trend was observed for surround suppression, a key property of macaque V1 that is thought to be mediated by lateral and feedback connections (Bair, Cavanaugh, & Movshon, 2003; Nassi, Lomber, & Born, 2013; Nurminen, Merlin, Bijanzadeh, Federer, & Angelucci, 2018). Surprisingly, we observe that even in purely feedforward hierarchical neural network candidate V1 models, single artificial neurons also exhibit suppression of responses from surrounding stimuli (Figure 2D and Supplementary Figure 1).

These results thus far qualitatively demonstrate that despite their much simpler architecture when compared to cortical circuits, single artificial neurons in certain hierarchical neural network models respond to visual stimuli similarly to single macaque V1 neurons. Furthermore, response properties of single artificial neurons in these hierarchical models depend on different aspects of the model commitment to biology (e.g. assumed field-of-view), and, in some cases, in unexpected ways.

### Distributions of single neuron properties in specific processing stages of certain hierarchical neural networks quantitatively approximate those in macaque V1

Because single artificial neurons in some of the V1 candidate models respond similarly to single neurons in macaque V1, we next sought to quantify the responses of these individual artificial neurons and compare them to those of many individual neurons in macaque V1. For example, it is possible that some V1 neuronal subpopulations are completely absent in some candidate V1 models or that the V1 model neuronal populations are biased towards some response types. Specifically, we compared the distributions of response properties in the V1 models with the respective empirical distributions measured in macaque V1. We focused on 22 single neuron response properties that we extracted from published V1 studies (Supplementary Table 1; Cavanaugh et al., 2002; De Valois, Albrecht, et al., 1982; De Valois, Yund, et al., 1982; Freeman et al., 2013; Ringach et al., 2002; P. H. Schiller et al., 1976; Ziemba et al., 2016) and replicated the corresponding experiments in each V1 candidate model (Figure 3A and Supplementary Figure 2). Each in silico experiment consisted of estimating an empirical model neuronal distribution of a random sample of artificial V1 neurons with the the empirical biological distribution of the same size (presumed random) sample reported in the corresponding neurophysiological experiment. This procedure was then repeated 1,000 times to estimate the uncertainty with respect to candidate V1 model neuronal sampling (methods). We considered the distributions of the following response properties: preferred orientation, circular variance (CV), orientation selectivity, orientation halfbandwidth, ratio of orthogonal and preferred responses (Orth./Pref.), ratio between CV and orientation half-bandwidth, difference between the Orth./Pref. and CV, peak SF, SF selectivity, SF bandwidth, grating summation field, surround diameter, surround suppression index, texture modulation index, absolute texture modulation index, F1/F0 ratio, texture selectivity, texture sparseness, texture variance ratio, maximum DC response, maximum texture response, and maximum noise response.

**Figure 3.**
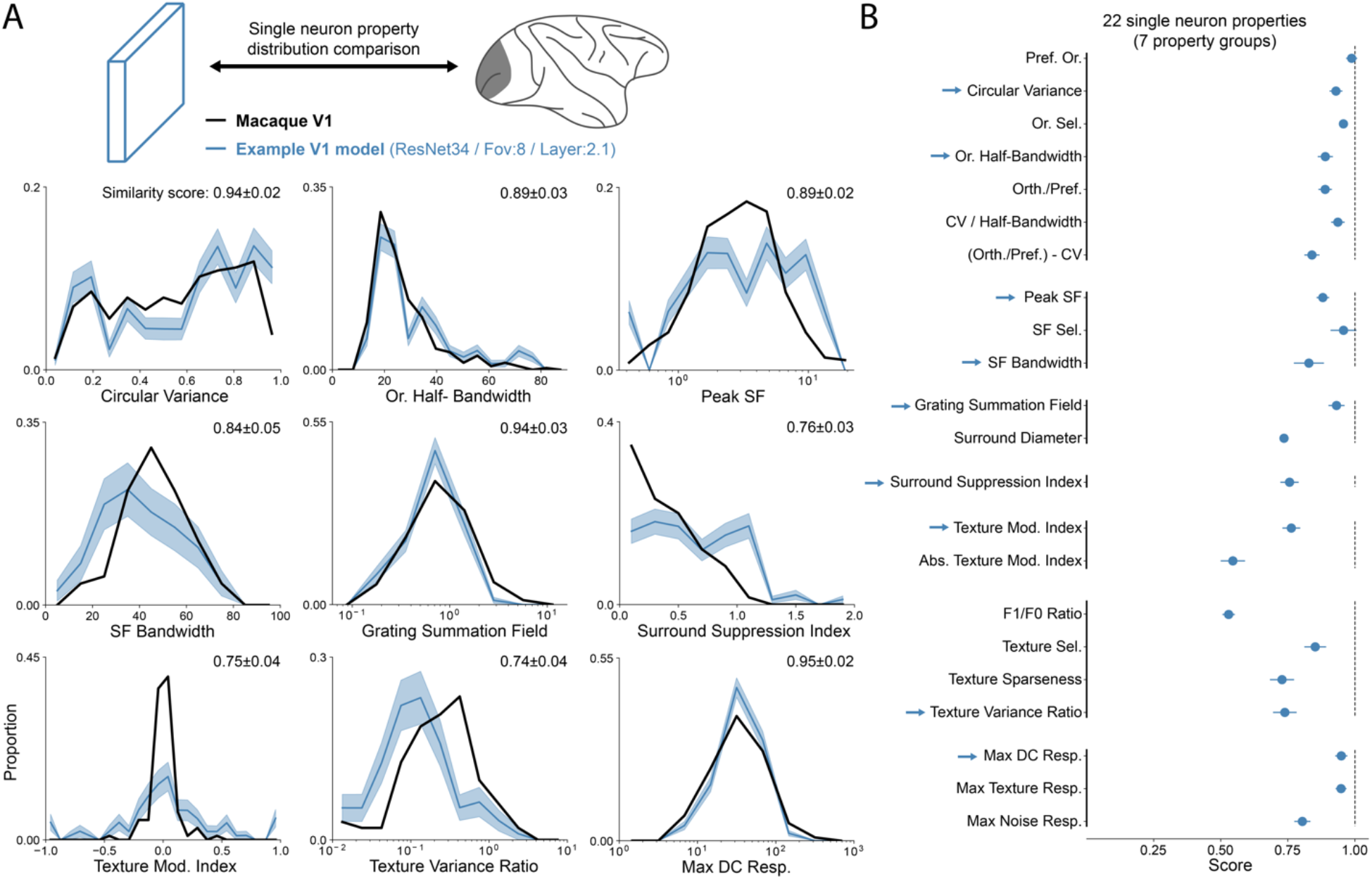
Distributions of single neuron response properties in a candidate V1 model approximately match those in macaque V1. **A.** Distributions of nine example response properties in macaque V1 (from published studies, black line) and a V1 model (ResNet34-FoV8-Layer2.1, same as in Figure 1). Model distributions are obtained by performing in silico experiments, thick blue line is the mean over 1,000 experiments and the shaded region is the SD. All the 22 response property distributions are shown in Supplementary Figure 2. Normalized similarity scores are shown in each plot at the top right corner. **B.** Similarity scores for the 22 single neuron response properties for the same V1 model (error bars represent mean and SD). Arrows indicate the response properties shown in **A.** Response properties are displayed in seven groups: orientation tuning, spatial frequency tuning, receptive-field size, surround modulation, texture modulation, response selectivity, and response magnitude.

We found that certain V1 models had distributions of response properties that closely approximated those reported in macaque V1 not only in their range but also their distributional shape (Figure 3A and Supplementary Figure 2). We defined a normalized distribution similarity score as (1 – *KS_c_^M–E^*)/(1 – *KS_c_^E–E^*), where *KS_c_^M–E^* is the ceiled (see methods) Kolmogorov-Smirnov (KS) distance between the empirical model distribution and the empirical biological distribution and *KS_c_^E–E^* is an estimate of the expected ceiled KS distance between different biology experiments. A low score means that the model distribution does not match the biological distribution while a score of 1 means that the model distribution is indistinguishable from the biological distribution considering experimental variability. We note that the similarity score ceiling is not limited by the choice of model. Instead, it depends on, and is thus limited by the biological sample size and number of bins of the empirical biological distribution: smaller number of neurons and bins tend to lead to lower ceilings (i.e. more uncertainty about the empirical biological distribution) and therefore decrease the range of scores for different models. To quantify how well a V1 model approximates macaque V1 at the single neuron level according to these response properties, we pooled the property scores in seven groups (Figure 3B): orientation tuning, spatial frequency tuning, receptive-field size, surround modulation, texture modulation, response selectivity, and response magnitude. Then, we averaged the scores within each group, and, finally, averaged the seven group scores, obtaining a V1 composite properties score (Supplementary Figure 3). This composite score serves as a summary of the match to the measures that were chosen for this study, weighted as outlined above.

Since the V1 composite properties score depends on the property distributions of the model’s individual V1 neurons, the composite score also depends on all of the factors outlined in the previous section: the choices of base model, FoV, and base model layer (Supplementary Figure 3). For V1 models derived from the same base model and FoV, similarity scores vary considerably with the choice of model layer to assign as V1 (Supplementary Figure 3A). In particular, the scores for different V1 properties show interesting dependencies on these choices that motivates future work: for example, receptive field size similarity is optimal for a subset of FoV and model layer combinations (as observed by a narrow band in Supplementary Figure 3B). While some V1 candidate models achieved very high scores for multiple response properties, no candidate V1 model tested here exactly matched macaque V1 in all the response properties (Figure 3B shows scores for the V1 model with highest V1 composite properties score).

In summary, we found that single artificial neurons of certain candidate V1 models contained within hierarchical neural network models exhibit responses similar to those of single neurons in macaque V1 and closely approximate V1 when considering distributions over many individual neurons. In spite of this, no model in the large pool of candidate V1 models analyzed (n=736) fully matched the macaque V1 along all the response properties.

### Different response property similarity scores provide complementary information about a model’s similarity to V1

Why is no single model able to match the distributions of all V1 response properties? One hypothesis is that there are some response properties that no model in the family of feedforward, ImageNet-trained ANN models considered here is able to approximate. An alternative hypothesis is that all of the biological V1 properties measured thus far are explained by this model family, but that they are found in different model layers rather than being expressed in a single population of putative V1 neurons. Distinguishing between these alternatives could guide future model architectural choices.

To disambiguate these two hypotheses, we first looked at the distributions of all the property scores over all the V1 models (Figure 4A). The distributions of scores for different properties varied considerably in their ranges: some properties, such as the preferred orientation and maximum DC responses had very high scores for most models, while others, such as the grating summation field and texture variance ratio, had broad distributions of scores. This is also illustrated by the large spread of score medians over the different properties, ranging from 0.41 for the surround diameter to 0.96 for the preferred orientation. Still, despite the large differences between the distributions of scores, for most properties, at least one of the candidate V1 models had a very high score. In particular, only three properties had a maximum score lower than 0.95 (surround suppression index, texture modulation index, and texture sparseness), and none lower than 0.9. In sum, the family of ANN-derived multi-stage models we considered is already capable of matching all of the 22 biological V1 response properties studied here – but no single model alone captures all of the response properties.

**Figure 4.**
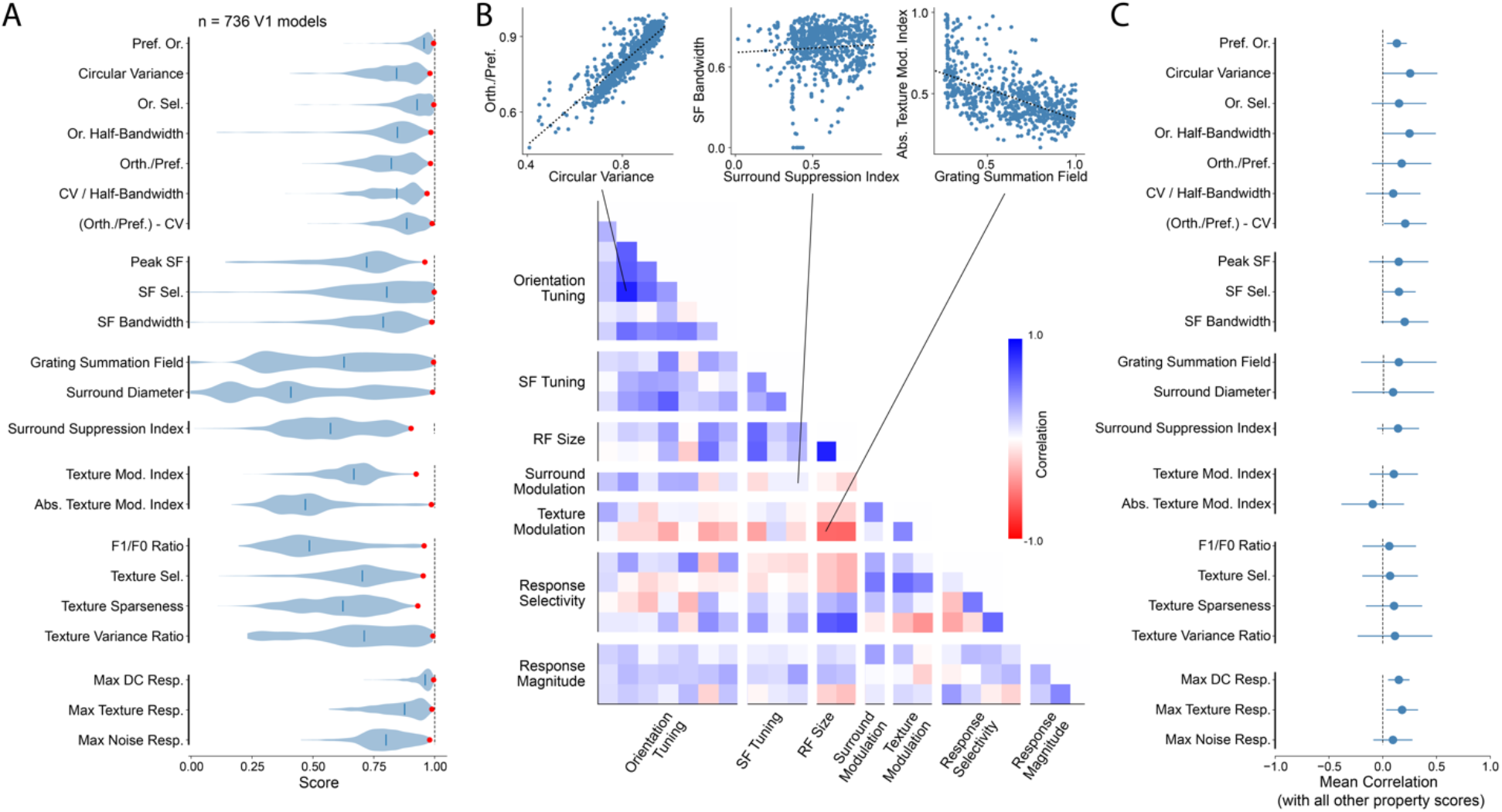
Single neuron response property similarity scores are on average weakly correlated across V1 candidate models. **A.** Violin plots show the distributions of similarity scores over 736 ANN V1 models for the 22 single neuron response properties. Thick blue lines indicate the median of each distribution and the red dots the maximum. There is significant variability in the individual property scores across models. **B.** Top, scatter plots comparing three pairs of different response property similarity scores. Left, similarity scores of circular variance and ratio between orthogonal and preferred orientation responses are positively correlated. Middle, similarity scores of surround suppression index and spatial frequency bandwidth are not correlated. Right, similarity scores of absolute texture modulation index and grating summation field are negatively correlated. Bottom, pair-wise correlations between the 22 single neuron response property scores grouped in the seven groups (correlations are calculated across all models). Lines connect to the corresponding scatter plots on top. **C.** Mean pair-wise correlations for each response property with all the others (errorbars represent mean and SD). On average single neuron response property scores are weakly correlated.

We then looked at the correlations between different property scores across all V1 candidate models. Correlations between scores of different properties showed great variability (Figure 4B). We found that some pairs of properties such as the ratio of orthogonal and preferred responses and circular variance were highly correlated over V1 models – models that tend to match one property also tend to match the other (Figure 4B top left, r=0.87, p-value=9.3E-232). Scores of other pairs such as the surround suppression index and spatial-frequency bandwidth are not correlated at all (Figure 4B top middle, r=0.04, p-value=0.29). Most interestingly, we found that some pairs of properties scores were anti-correlated (Figure 4B top right), e.g. grating summation field and absolute texture modulation index (r=-0.59, p-value=3.9E-70). That is, V1 models that capture one property, tend to do worse on the other property. We also found that scores of properties that belong to the same group, i.e., that relate to similarly named phenomena, were significantly more correlated than scores of properties of different groups (r_same_= 0.33±0.28 vs r_different_= 0.12±0.23; t-test, t=4.04, p-value=2.2E-4). When considering all the properties, scores were on average weakly correlated (Figure 4C).

Next, we analyzed whether the property scores could be explained by simple model parameters. As previously mentioned, within the same base model, some property scores depend on model parameters such as the FoV and layer depth (Supplementary Figure 3). When we consider all the models, we observed some interesting relationships between property scores and model parameters that were not always aligned (Supplementary Figure 4A). For example, both the receptive field size and the texture modulation property scores vary with the model’s theoretical receptive field and layer depth but were optimal for different values (Supplementary Figure 4A). We then performed a sequential ANOVA to identify which model parameters contributed the most to explain variance in the property scores and quantify how much of the variance in scores can be attributed to the model parameters (Supplementary Figure 4B; model parameters considered were the model total depth, FoV, layer depth, theoretical receptive field, layer type, and number of neurons). Theoretical receptive field was the most important model parameter in explaining the variance in scores of four property groups (orientation tuning, spatial frequency tuning, receptive field size, and response magnitude) as well as in the V1 composite properties. On the other hand, layer depth was the model parameter that explained the most variance for surround modulation, texture modulation and response selectivity properties. In total, the six model parameters accounted for 67.3% of the variance in the V1 composite properties scores.

These results show that within the set of V1 candidate models analyzed here, there is at least one model that approximates each of the macaque V1 property distributions reasonably well (score over 0.9). However, since scores of different properties are on average weakly correlated and some pairs of property scores are, in fact, anti-correlated, no model was able to simultaneously match all of the V1 empirical distributions. This suggests that different properties reflect different aspects of model similarity to V1, which are not necessarily aligned, and that there may exist constraints in the model architecture (e.g. feedforward only except for the CORnet-S architecture), limiting the ability to fully approximate biological V1. Finally, a large fraction of the variance in the scores can be explained by simple model parameters such as the theoretical receptive field and layer depth.

### Single neuron property similarity scores correlate with similarity scores derived from standard neural predictivity metrics

Conventional model evaluation methods deployed over the last several years fit a map between model neurons and individual biological neurons and then score the ability of the model to predict the responses of each mapped biological neuron to new (held out) stimuli such as complex images. How do the V1 property scores described here compare to these conventional methods that evaluate the model’s similarity to V1 (Cadena et al., 2019; Dapello et al., 2020; Schrimpf et al., 2018)? To address this question, we calculated for each V1 candidate model how well it explained stimulus driven responses of V1 neurons using a conventional neural predictivity methodology based on the partial least square regression (PLS) model mapping method (Helland, 2006; Schrimpf et al., 2018; Yamins et al., 2014). We used a neuronal dataset containing extracellular recordings from 102 single-units while presenting naturalistic textures and noise images which had been originally published in a study analyzing texture modulation in macaque V1 and V2 (Freeman et al., 2013). The dataset consisted of stimulus-evoked responses to 315 images (20 repetitions and averaged over 150ms). For each model, biological neurons were mapped to the V1 model neuronal population linearly using a PLS regression of the model with 25 components. Model predictions were evaluated using a 10-fold crossvalidation strategy. V1 explained variance was then normalized by the neuronal internal consistency to arrive at the specific V1 explained variance benchmark that we consider next.

Across all V1 models, we found that the neuronal explained variance benchmark was strongly correlated with the V1 composite properties score outlined above (Figure 5A, r=0.61, p-value=1.8e-76). On average, individual property groups were also correlated with the V1 explained variance benchmark, though there was considerable variability across groups (Figure 5B,C; r=0.31±0.09, mean and SD). Increasing the number of property groups averaged gradually improves the alignment of the V1 component property scores with explained variance under this dataset (Figure 5B). This correlation was not exclusive to V1 explained variance using PLS regression, since it was present when using other neural predictivity methods on this same neural dataset. In particular, response property distribution similarity scores were also correlated with explained variance using single neuron mapping (choosing the single neuron in the model that best predicts a single macaque V1 neuron; r=0.50, p-value=9.4E-49) (Arend et al., 2018), and with representational similarity metrics such as representational dissimilarity matrix (RDM; r=0.62, p-value=6.9E-79) (Kriegeskorte et al., 2008), and center kernel alignment (CKA; r=0.23, p-value=5.7E-10) (Kornblith, Norouzi, Lee, & Hinton, 2019) that do not involve fitting model features. Considering all the different metrics tested, we found that spatial frequency tuning and response selectivity were the V1 component property scores that were most correlated with V1 explained variance and representational similarity on this particular V1 dataset (Figure 5C). Finally, the alignment between the V1 composite properties scores and the V1 explained variance was not an artifact of the way that the individual property scores were first averaged in groups (Supplementary Figure 5). When directly calculating the mean of the 22 individual property scores, the correlation with V1 explained variance persisted (r=0.56, p-value=2.7E-62), even when removing the seven response properties that overlap with the neuronal dataset used for determining the explained variance (r=0.42, p-value=1.0E-32, methods).

**Figure 5.**
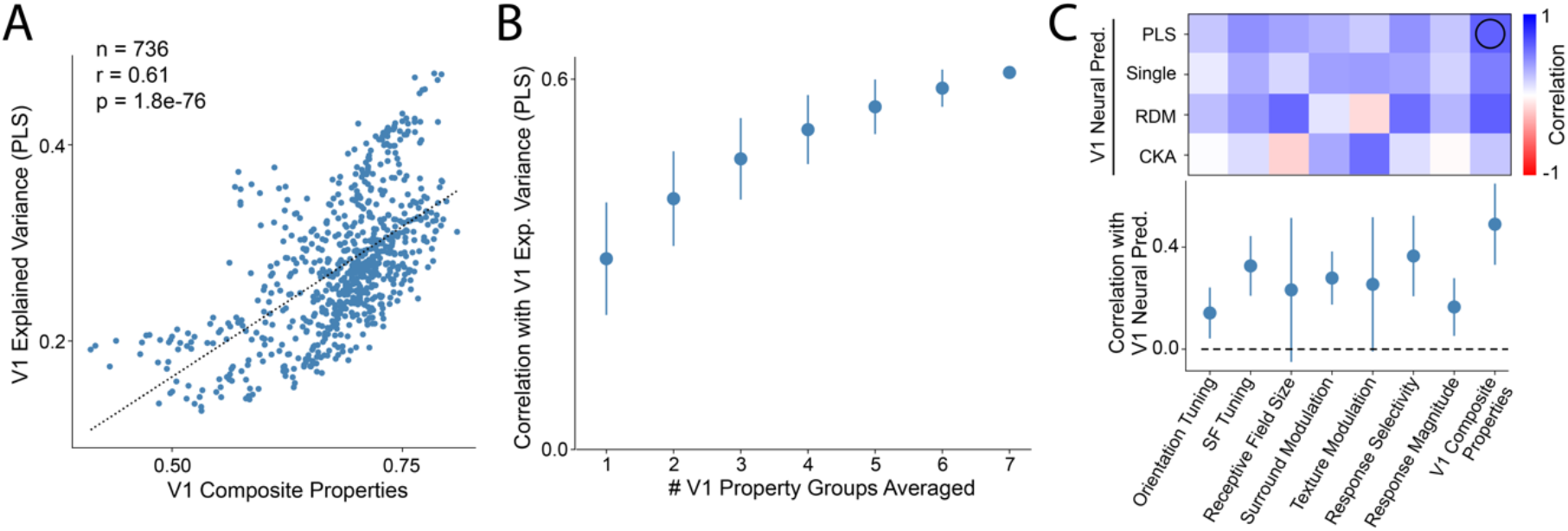
Single neuron response property similarity scores correlate with the V1 model’s ability to predict neuronal responses using standard measures (cross-validated explained variance and representational similarity analysis). **A.** Comparison of model’s ability to explain variance in macaque V1 responses in a neuronal dataset (Freeman, Ziemba et al 2013) using PLS regression and V1 composite properties scores (across 736 V1 models). Model’s cross-validated explained variance is positively correlated with the V1 composite properties scores. **B.** Averaging over an increasingly higher number of property group scores improves correlation with V1 explained variance. **C.** Top, correlation of V1 property groups and composite properties scores with different V1 neural predictivity metrics (explained variance with PLS regression and single neuron mapping, and representational similarity analysis with RDM and CKA). Open black circle indicates the correlation shown in **A**. Bottom, same as above but showing the mean and SD across the different explained variance metrics.

In sum, we found that the V1 property scores are partially aligned with more conventional methods for evaluating a candidate V1 model’s similarity to V1. While this result is not entirely surprising, it serves as an important sanity check, showing that there is signal in the response property distributions similarity scores for benchmarking models in how well they explain V1. In addition, this result can be interpreted as at least a partial validation of regression-based methods for evaluating neural predictivity of ANN-based models, in spite of their reliance on a fitting step (see Discussion).

### Hierarchical models that have more brain-like V1 stages are more aligned with primate object recognition behavior

While accurately modeling primate V1 is an important goal in and of itself, our larger goal is to do this in the service of multi-scale, multi-stage models of the ventral visual stream and visually-driven behaviors. Thus, we next asked: do multi-stage artificial neural network models that better align with biological V1 also tend to better align with biological behavior? Building on prior work, we here focused on primate core object recognition behavior. Specifically, when assessed via batteries of core object recognition tasks, humans and monkeys show highly aligned difficulty and confusion patterns at the object- and image-level (Rajalingham et al., 2015; Rajalingham et al., 2018). That is, humans and monkeys not only show similar levels of accuracy in a visual categorization task, but they also reliably show the same patterns of successes and failures when scored at the grain of object categories (pooling over subjects and images of the same category) or at the grain of individual images (pooling over subjects), and those reliable patterns can thus be used to assess the biological fidelity of any image-computable model at the behavioral level. Indeed, while some hierarchical neural network models accurately match typical primate patterns of object confusion, they do not yet match those patterns at the image-level (Rajalingham et al., 2018) and some models match better than others (Schrimpf et al., 2018 and accompanying website). Thus, we here asked if hierarchical models with intermediate layers that better match V1, i.e. more brain-like V1-layers, also better match human (and monkey) core object recognition patterns of behavior using these same prior benchmarks.

Each hierarchical model with a specific FoV was here taken as a candidate multi-scale, multi-stage neural network model of the ventral stream and its resultant behavior. For each of these candidate ventral stream models (n=52, 13 base models with 4 different FoVs), we chose the layer that best approximated macaque V1 according to the layer’s single neuron composite properties score (above). We then performed psychophysical experiments on each model to evaluate how well the overall model aligned with human image-level behavior in classifying a set of naturalistic synthetic images using the same benchmarks and methods described in prior work (Rajalingham et al., 2018): Images consisted of objects belonging to 24 categories (10 images per category, 240 in total) displayed at different positions, orientations, and sizes, and overlayed on a random natural background. For each hierarchical model, we used standard methods of producing object recognition behavior from each model (Rajalingham et al., 2018). Specifically, we trained a behavioral decoder to classify the images using the activations of the model’s penultimate layer (last layer before the 1000-way class probability layer trained on ImageNet). The behavioral decoder was trained on a separate set of 2160 images of the same object categories.

We found that match (aka “consistency”) of the hierarchical neural network models with human image-level behavioral patterns was strongly correlated with the model’s V1 composite properties score of its most V1-like internal layer (Figure 6A, r=0.79, p-value=5.0e-12). Reducing the number of V1 property groups that are included (averaged) in the V1 properties composite score reduced the correlation between model V1 match scores and model behavioral consistency with humans (Figure 6B). This suggests that all V1 properties measured here are at least partially important to understanding V1’s role in supporting recognition behavior. However, we also found considerable variability between the correlation of single response property scores and behavioral consistency under this benchmark, with response selectivity showing little to no correlation, and orientation tuning being anti-correlated (Figure 6C). Despite object-level behavioral consistencies being considerably higher than image-level consistencies, the alignment between V1 property scores and behavioral consistency was very similar for all the different behavioral metrics (Figure 6C). Finally, the V1 similarity on the neuronal dataset previously described, as measured by the explained variance and representational similarity analysis, was considerably less correlated with behavioral consistency than the V1 composite properties scores (Figure 6C).

**Figure 6.**
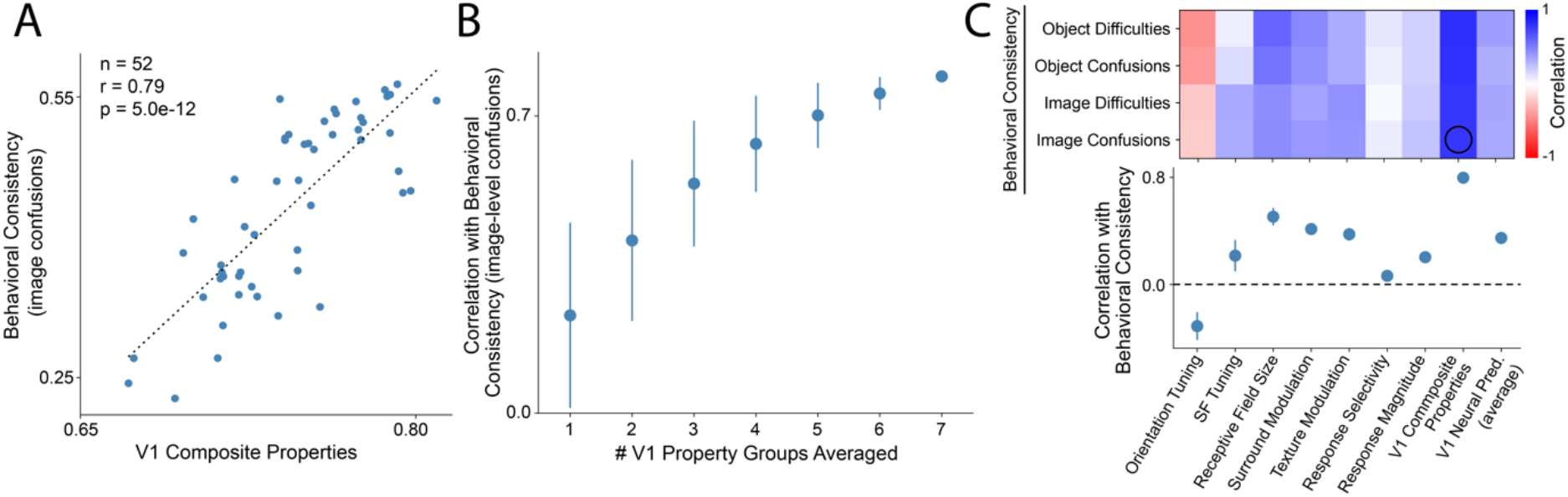
Hierarchical models with internal processing stages (layers) that better match macaque V1 at the single neuron level have output behavioral patterns that are better matched to human behavioral patterns. **A.** Comparison of human object recognition behavioral consistency and V1 composite properties scores for 52 hierarchical models. Behavioral consistency represents the alignment between the models’ output classification (i.e. model “behavior”) and humans performing a two-alternative forced choice object recognition task at the image-level (I2n metric, image-level normalized confusion patterns, see (Rajalingham et al., 2018; Schrimpf et al., 2018)). For each hierarchical neural network, the internal model layer with the highest V1 composite properties score was chosen. **B.** Increasing the number of V1 response property groups considered in the overall V1 score improves the correlation with behavioral consistency (the plot in A shows the result when all seven property groups are included). **C.** Top, correlation of V1 property group scores, V1 composite properties scores and V1 neural predictivity (average of explained variance and representational similarity analysis metrics) with behavioral consistency metrics with varying granularity: object difficulties, object confusion patterns, image normalized difficulties, and image normalized confusion patterns. Open black circle indicates the correlation shown in **A**. Bottom, same as above but showing the mean and SD across the different behavioral consistency metrics.

In sum, we found an empirical, multi-scale linkage between single neuron response properties and object recognition behavior: multi-stage neural network models with an internal V1 layer that better matches the distribution of individual V1 biological neuronal properties tends to also better match biological behavior.

### How do the candidate V1 models tested here compare with classical models of V1?

As previously mentioned, no V1 model derived from the family of ventral stream models considered here was able to completely match macaque V1 along all the single neuron response properties considered. We wondered whether a classical neuroscientific model of V1 might do better, given that such models were constructed largely guided by the kinds of empirical results used here. While those classical models are limited in that they do not bridge all the way to behavior, their match to V1 is nonetheless an important reference in determining future ventral stream modeling efforts. Answering how good the classical neuroscientific model is turns out to be non-trivial as there is no standard, agreed-upon classical neuroscientific model. For example, Cadena et al. showed that a task-optimized ANN outperformed one variant of the classical linear-nonlinear model of V1 based on a Gabor filter bank (GFB) followed by simple- and complex-cell nonlinearities in predicting responses in macaque V1 (Cadena et al., 2019). However, Dapello, Marques et al. more recently showed that constraining the GFB parameters with empirical data (i.e. a different variant of the classical model) substantially improves its ability to explain V1 response variance (Dapello et al., 2020).

To answer our original reference question (above) and to help clarify the current state of the art in image-computable V1 models, we implemented both of these V1 classical models: a data-constrained classical V1 model consisting of a GFB (J. P. Jones & Palmer, 1987), simple- and complex-cell (Adelson & Bergen, 1985) nonlinearities and a simple divisive normalization stage (Carandini, Heeger, & Movshon, 1997; Heeger, Simoncelli, & Movshon, 1996) (classical V1 model variant 1; see Methods), and the GFB model used in Cadena et al. (classical V1 model variant 2). We then compared both of these classical model variants with the candidate V1 models derived from hierarchical ANNs considered in this study (above). Using the V1 properties scores developed here, we found that the classical V1 model variant 1 outperformed all the ANN-derived V1 models by a wide margin, but that many of the ANN-derived models outperformed the classical V1 model variant 2 (Figure 7A,C; V1 composite properties scores: classical V1 model variant 1 0.90±0.03, classical V1 model variant 2 0.70±0.03, best V1 model based on hierarchical ANNs 0.81±0.03).

**Figure 7.**
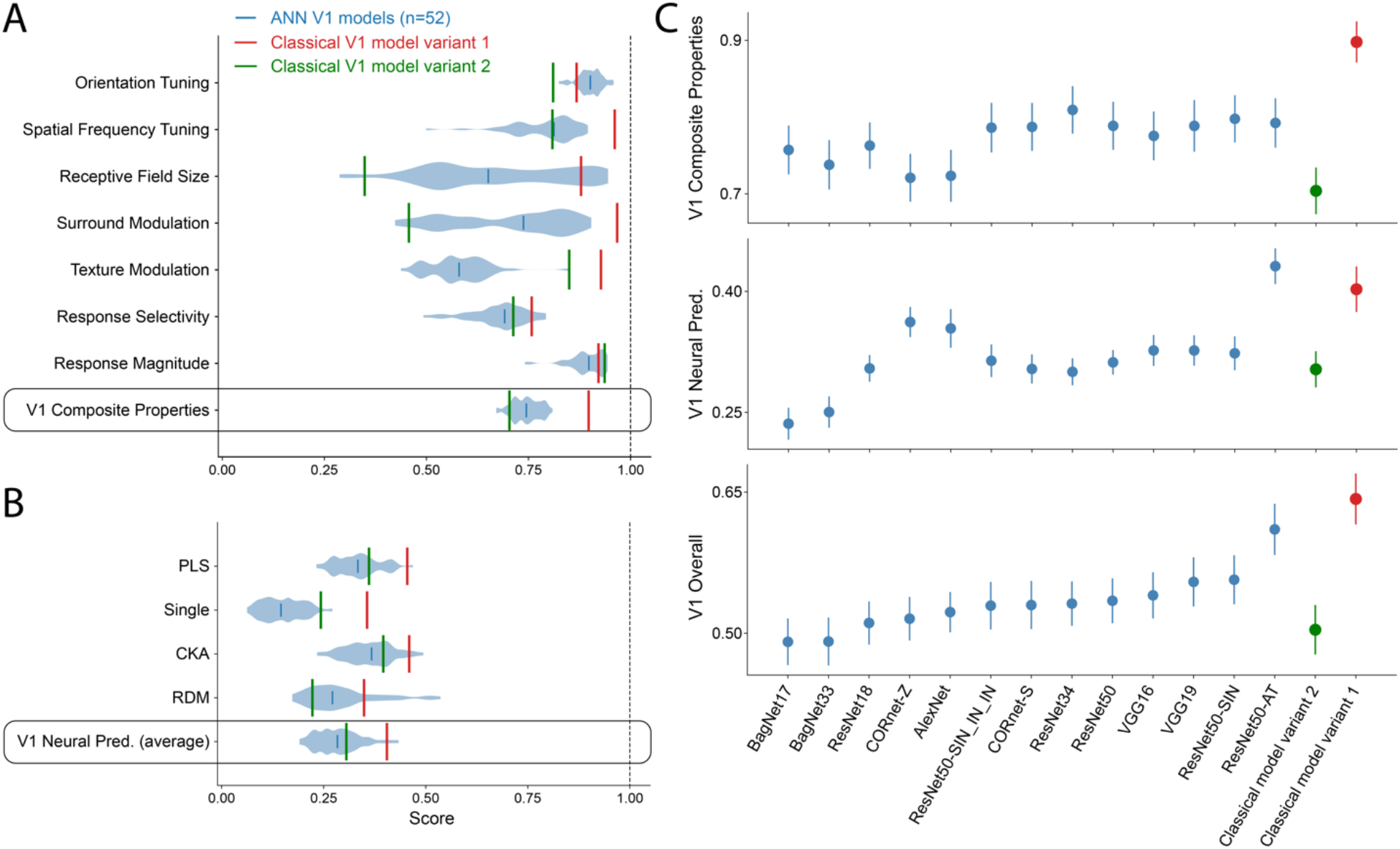
An optimized classical neuroscientific V1 model outperforms V1 models based on hierarchical neural network models in approximating macaque V1. **A.** Violin plots represent distributions of scores for the seven groups of V1 properties as well as the V1 composite properties for the most V1-like layer in 52 hierarchical models (13 base models with 4 FoVs). Green line represents the GFB model used in Cadena et al 2019 as the classical V1 model control. Red line represents an optimized neuroscientific model of V1 with a GFB, simple- and complex-cell nonlinearities, and divisive normalization stage. **B.** Same as in **A** but for V1 neuronal predictivity using different metrics (PLS regression, single neuron mapping, CKA, and RDM) and the average of all metrics. **C.** V1 composite properties score (top), average neuronal predictivity (middle), and V1 overall score (bottom) for the best combination of layer and FoV for each base hierarchical model and the two classical V1 models. V1 models based on hierarchical ANNs are sorted according to their V1 overall score. An optimized classical V1 model outperforms all hierarchical models on the V1 composite properties scores and overall and is only surpassed by an adversarially-trained model in neuronal predictivity. The adversarially-trained ResNet50 outperformed all other models in V1 neuronal predictivity.

Interestingly, when using standard V1 neural predictivity metrics (explained variance and representational similarity analysis) on the neuronal dataset described before, we found that a V1 model derived from an adversarially-trained hierarchical ANN (Engstrom, Ilyas, Santurkar, & Tsipras, 2019; Madry, Makelov, Schmidt, Tsipras, & Vladu, 2019) was better than all the other ANNs (Figure 7B,C; average V1 neural predictivity: adversarially-trained ANN 0.43±0.02, classical V1 model variant 1 0.40±0.03, classical V1 model variant 2 0.30±0.02, best standard ImageNet-trained ANN-based model 0.36±0.02). This result had already been reported by some of the authors (Dapello et al., 2020) but here we extend to other metrics.

In sum, using the V1 composite properties score, the classical V1 model variant 1 is a better match to macaque V1 than all the ANN-derived V1 models explored in this study. The same holds true even when averaging both the V1 composite properties scores with the V1 neural predictivity (Figure 7C, bottom). However, none of the models we tested were able to fully approximate biological V1. Together, these results suggest that there is still considerable room to improve current models of macaque V1, which will likely lead to better ventral stream models of core object recognition.

## Discussion

In this work, we evaluated whether hierarchical neural networks of artificial neurons are accurate, multi-scale, multi-stage models of the first stage of cortical processing of the primate ventral visual stream. The long-term goal of such work is to use such models to bridge between the properties of individual cells all the way to visually driven behaviors such as object recognition. While it was known that some of the models we tested are moderately accurate in predicting neuronal responses along the ventral stream areas using regression methods and representational similarity analysis, how their single neurons relate to single neurons in the brain had not been studied before, to the best of our knowledge.

Specifically, our goals in this study were: (1) to build end-to-end neural network models of the ventral stream (using known hierarchical neural network architectures, but here exploring and specifying critical open parameters such as model FoV), (2) to evaluate whether single neurons in V1 levels of those new, specific brain models were functionally similar to single neurons in macaque V1; (3) to test whether the distributions of single neuron response properties in the V1-layers of these models matched those in macaque V1 for a range of previously reported core biological response properties; and (4) to ask if hierarchical models with better V1 layers (i.e. more similar to macaque V1 at the single neuron level) were also more similar to primate behavior at their output (i.e. behavioral) level.

Our results show that single neurons in V1-layers of hierarchical models of the primate ventral visual stream have response characteristics that are surprisingly similar to their biological counterparts, and also that the distributions of response properties in some models’ V1-layers approximately match those in macaque V1. Furthermore, we observed that hierarchical models with more brain-like V1-layers, were also more aligned with human object recognition behavior (at their output layers). Together with prior work, these results suggest that the hierarchical neural network models that we built here are reasonably accurate, multi-scale models of the primate ventral visual stream and its support of object recognition behavior. We have made the specific leading models we found here publicly available for community use and further exploration. Nevertheless, these results also show that none of these models are perfect, and they provide pointers to future modeling improvement avenues.

### Multi-scale, hierarchical models of the primate ventral stream

Certain hierarchical artificial neural networks have achieved unparalleled accuracy in predicting visual neuronal responses in low- (Cadena et al., 2019) and high-level (Yamins et al., 2014) visual cortical areas, as well as object recognition behavior (Rajalingham et al., 2018), making these specific models the current best scientific hypotheses of the neural processing mechanisms at work along the primate ventral stream (Schrimpf et al., 2018). However, these models are far from perfect. For instance, they cannot completely account for all the explainable response variance in the neural and behavioral data (Schrimpf et al., 2018). Furthermore, these models tend to be largely feedforward, making it hard for them to model the recurrent processing dynamics in the ventral stream; though we and others have recently started to address this (Kietzmann, Spoerer, Sörensen, Cichy, & Hauk, 2019; Kubilius et al., 2019; Nayebi et al., 2018; Tang et al., 2018).

In this study, we focused on another critical aspect that the leading hierarchical ANNs still lack: if any such model claims to be a causal, multi-scale, multi-stage, neurally-mechanistic model of the primate ventral visual stream it must have key parameters fully specified (e.g. FoV) and it must have alignment with cortical areas at the level of individual neurons. Specifically, past studies using multi-stage ANN models to explain single neuronal responses in primate ventral stream areas have used the model neurons as an encoding basis (Cadena et al., 2019; Schrimpf et al., 2018; Yamins et al., 2014) or with representational similarity analysis (Cadieu et al., 2014; Güçlü & van Gerven, 2015; Khaligh-Razavi & Kriegeskorte, 2014). While the first approach accurately fits and predicts individual biological neuronal responses, it does so by combining the activity of thousands of model neurons and it also implicitly allows key model parameters to be left unspecified. Similarly, the latter approach compares the representational spaces between a neuronal population in the model and a neuronal population in the brain. In both cases, single neurons in the model are not explicitly mapped to individual neurons in ventral stream areas. Here, we take first steps towards hierarchical models of the primate ventral stream that are truly multi-scale, multi-stage and thus can bridge between properties of individual cells in intermediate model levels to visually-guided behavior.

Our approach consisted in hypothesizing a one-to-one mapping at the level of single neurons between macaque V1 and a particular choice of processing stage in each hierarchical model with a committed FoV which we refer to as a candidate V1 model. Then, by performing a series of in silico experiments replicating neurophysiological studies, we characterized single neuron responses in the model V1-layer and compared them to those in the macaque V1. This approach presents multiple advantages when compared to existing methods for evaluating V1 similarity. The first, and most obvious one, is the one-to-one single neuron mapping which ensures the alignment at the level of single neurons between the candidate V1 model and the corresponding cortical area. This, in turn, dispenses the standard regression fitting procedure, which can give rise to situations of under- and over-fitting depending on the availability and variability of the data. Another important advantage of the approach used here is the improved interpretability of the response property distribution similarity scores. Conventional methods for evaluating a model’s match to neuronal data only provide an answer to the question “How much variance does the model explain?”. Our approach, on the other hand, gives a more detailed description of “Which aspects of the neuronal responses are explained by the V1 model and which aspects are not?” In addition, it does so in a language familiar to the visual neuroscience community by relying on single neuron response properties extensively used in V1 neurophysiology. Finally, the proposed methodology builds on *published* primate V1 studies without requiring additional recordings, meaning that the approach presented here can be further extended to include other aspects of V1 processing without requiring new biological experiments. Other single neuron response properties such as those related to color tuning (Horwitz & Hass, 2012), figure-ground modulation (Lamme, 1995), and border ownership (Zhou, Friedman, von R, & von der Heydt, 2000) are viable properties to operationalize into quantitative benchmarks to give an even more complete evaluation of the model’s similarity to V1 at the level of single neurons.

### Modeling macaque V1 with hierarchical neural network models at the level of single neurons

We here report that some hierarchical neural network models provide moderately accurate models of macaque V1 at the level of single neurons. This result extends prior work that used similar models to predict neuronal responses in macaque V1 using regression-based methods (Cadena et al., 2019; Dapello et al., 2020; Schrimpf et al., 2018). However, we report that no model was able to simultaneously match all the 22 single neuron response property distributions (highest V1 composite properties score was 0.81±0.03). This was not due to some individual response properties being impossible to match by this model family since there was always at least one model that approximately matched each of the V1 single neuron response property distributions (similarity score larger than 0.9). Instead, the reason for the failure to match all of the response properties is likely due to the limitations of the model architectures we considered. Scores for some response properties varied with particular model properties and in several cases in non-optimal ways.

Not surprisingly, we found that V1 similarity at the level of single neurons was somewhat aligned with V1 neuronal predictivity using conventional approaches (explained variance and representational similarity analysis). Still, this alignment was not perfect suggesting that each individual comparison benchmark reflects only a particular, and incomplete, measure of a given model’s match to biological V1. This is true for each of the single neuron property distribution similarity scores, which only evaluate V1 similarity in a very specific phenomenon, as well as for the V1 neuronal predictivity which is limited by the type of stimuli and size of the dataset. This observation reinforces the idea that, to best evaluate the ability of a model to explain brain processing, one should consider multiple and varied benchmarks in an integrative manner (Schrimpf et al., 2018, 2020).

Unlike Cadena et al. 2019, we found that V1 models derived from hierarchical ANNs underperformed in explaining V1 neuronal phenomena when compared to an optimized classical neuroscientific model. This was true for both the single neuron property distribution similarity scores and the V1 neuronal predictivity (except for the adversarially-trained model). While there are several differences in the neuronal datasets and the fitting procedures used in the two studies, we believe that the main reason for this apparent discrepancy in results is due to differences in the classical V1 models used. Indeed, when we used an off-the-shelf classical V1 model identical to the one used in the Cadena et al. study (here called “variant 2”), we qualitatively replicated their result in that we also found that many ANN-derived V1 models matched the biological V1 better than that classical V1 model. However, we also found that a better optimized classical V1 model (here called “variant 1”) that outperforms (on average) all of the tested ANN-derived models. This has important implications since it suggests that current hierarchical ANN architectures are not yet the best model class for approximating macaque V1.

How can we further improve models of V1? Our results suggest several possibilities. First, by tweaking the architectural parameters of hierarchical neural networks, such as the kernel sizes of the convolutions, it may be possible to remove the observed sub-optimal interactions and, thus, improve V1 similarity. In addition, we can expand the model space by including new circuit motifs inspired by neurobiology, such as local recurrence. The implementation of these types of architectural changes could potentially be guided by the V1 similarity scores described here. Another alternative is exploring other task-optimization procedures for training the model weights. Similarly to another recent study (Dapello et al., 2020), we found that adversarially-trained models had the highest V1 neuronal predictivity using conventional metrics. Exploring other types of data augmentation during training may result in even stronger V1 similarity. Finally, the higher V1 similarity scores of the classical neuroscientific model suggests that there may be opportunities in merging that type of mathematically elegant model with task-optimized neural networks. Such an approach has been successfully used to improve the adversarial robustness of neural network models for object recognition (Dapello et al., 2020). However, we must remain wary of the possibility that the space of firing-rate based hierarchical architectures considered here may not allow to completely approximate macaque V1.

### Bridging from single neuron properties to behavioral phenomena

Prior to this work, it might have been argued that modeling individual V1 neurons is a low level problem for neuroscientists and modeling object recognition behavior is a high level problem for cognitive scientists. Here we empirically demonstrate – for the first time -- that those two problems are closely intertwined and that a unified systems modeling approach produces gains in both. Building such a bridge is a longstanding goal of systems neuroscience, and models of the type tested here provide a path to reach that goal.

By evaluating the ability of hierarchical neural network models to match macaque V1 at the single neuron level, we were able to produce models that offer a bridge from the level of single neuron properties in the first visual cortical area all the way to object recognition behavioral phenomena. The alignment of V1 similarity with behavioral consistency in these hierarchical models suggests that object recognition behavior is derived, at least in part, from low-level visual processing in V1. This is in agreement with the recent result showing that V1 similarity also correlates with robustness to image perturbations (Dapello et al., 2020). Together, these results all point towards the same conclusion: that working to build better models of low-level visual processing has tangible payoffs in improving models of visual behavior.

Going forward, we believe that the approach used here will lead to a deeper understanding of the neural processing along the ventral visual stream and its support of visual object recognition and other visually driven behaviors. By building even more accurate, multi-scale, multi-stage models of the primate ventral stream, we will begin to uncover how specific cellular mechanisms at work along the multiple ventral stream areas contribute to different aspects of visual intelligence.

## Acknowledgments

We thank J. Anthony Movshon and Corey M. Ziemba for access to the V1 neural dataset, and Joel Dapello for adding some ANN models used in this study to the Brain-Score platform. This work was supported by the PhRMA Foundation Postdoctoral Fellowship in Informatics (T.M), the Semiconductor Research Corporation (SRC) and DARPA (M.S., J.J.D.), the Massachusetts Institute of Technology Shoemaker Fellowship (M.S.), Office of Naval Research grant MURI-114407 (J.J.D.), the Simons Foundation grant SCGB-542965 (J.J.D.), the MIT-IBM Watson AI Lab grant W1771646 (J.J.D.)

## Author Contributions

T.M. and J.J.D. designed the study. T.M., M.S., and J.J.D. designed the analysis. T.M. and M.S. implemented the analysis. T.M. developed the classical V1 model. T.M. and J.J.D. wrote the paper.

## Methods

### Hierarchical neural network models of the primate ventral visual stream

In this study we considered 13 pre-trained artificial neural networks (ANNs) to build multiscale, hierarchical models of the primate ventral visual stream. All models were accessed via the Brain Score platform (Schrimpf et al., 2018). The models considered were the following: AlexNet (Krizhevsky et al., 2012), VGG16 and VGG19 (Simonyan & Zisserman, 2015), CORnet-Z and CORnet-S (Kubilius et al., 2019, 2018), ResNet18, ResNet34 and ResNet50 (He et al., 2016), bagnet17 and bagnet33 (Brendel & Bethge, 2019), ResNet50-SIN and ResNet50-SIN_IN_IN, which are trained on a stylized ImageNet dataset (Geirhos et al., 2019), and an adversarially-trained ResNet50 with a ∥*δ*∥_∞_ = 4/255 constraint (Engstrom et al., 2019). All models were implemented with Pytorch (Paszke et al., 2019) except the VGG models which used Keras (Chollet, 2015). For each ANN base model, we built multiple V1 models using a three-step commitment as described in the Results section for a total of 736 V1 models.

### Empirical macaque V1 single neuron property distributions

To evaluate the ability of hierarchical neural network models of explaining macaque V1 at the single neuron level, we extracted from the literature the empirical distributions of 22 single neuron response properties in macaque V1 (Supplementary Table 1). Since some response properties depend on eccentricity, we chose distributions that corresponded to foveal neurons (eccentricities less than 5deg). In most cases, the distributions were directly taken from the paper figures using an assistance digital tool (WebPlotDigitizer). The distributions were rounded to integers and normalized to ensure the same total number of neurons reported in the paper. The single neuron response properties from Ringach et al. 2002 are publicly available online (http://ringachlab.net). The single neuron response properties from Freeman, Ziemba et al. 2013 and Ziemba, Freeman et al. 2016 were calculated from the neuronal responses generously provided by the authors.

### In silico neurophysiology experiments

To calculate the V1 model distributions for all the single neuron response properties, we performed in silico neurophysiology experiments that attempted to approximate the biological experiments carried out in the V1 studies. However, to facilitate their implementation, we made a methodological simplification that does not alter our results. Since all the layers of the ANNs considered here were convolutional layers, neurons respond identically at all the locations of the visual space. Due to this, instead of randomly sampling neurons with receptive fields (RF) spread along the whole visual space, we fixed a single location to record where we centered the stimuli (deviated from the center of gaze by 0.5deg on each axis resulting at an eccentricity of 0.7deg).

We first estimated the functional receptive field (RF) for all the neurons in the V1 model by presenting small gratings (0.33deg diameter, 3cpd spatial frequency, four orientations and two opposing phases) in a grid with a spacing of 0.25deg, and averaging the responses for each position. We then selected all the neurons that had their RF centers aligned with the stimulus location (within 0.15deg).

Depending on the single neuron response property, we presented the respective stimulus set. In total we used four different stimulus sets: gratings for orientation tuning (8 phases, 12 orientations, 4 spatial frequencies, 3 diameters per spatial frequency), gratings for spatial frequency tuning (8 phases, 6 orientations, 22 spatial frequencies, 1 diameter), gratings for size tuning (8 phases, 6 orientations, 4 spatial frequencies, 12 diameters), naturalistic textures and noise images (for both types 15 texture families and 15 samples per family). In addition, we used a homogeneous gray stimulus for obtaining the baseline responses. For each model, to convert its artificial neurons activations to neuronal firing rates, we calculated the affine transformation such that the median of the baseline responses and the median of the maximum DC responses in the orientation tuning stimulus set matches those in V1 (median of baseline responses = 0.82 spikes per second, sps; median of maximum DC responses = 33.8 sps, Ringach et al. 2002). We then applied the corresponding affine transformation for each model activations to obtain the firing rates. For each study, we selected responsive neurons based on a responsiveness criterion that emulates the one used in experimental characterization. Response properties for each neuron were then calculated from the responses to the corresponding stimuli following the methodology used in the original study. In some cases, the neuronal responses were baseline subtracted prior to calculating the property (Supplementary Table 1).

We sampled the same number of artificial neurons as those sampled in the empirical biological distribution of the corresponding experimental study (Supplementary Table 1). For some properties the number of neurons sampled in the models does not correspond to the number of neurons included in the neurobiological empirical distribution. For example, 87 neurons were considered for spatial frequency tuning in the Schiller et al study, but of those only 73 were SF selective and therefore included in the SF bandwidth distribution in the study. When performing these experiments in the model we sampled the same number of the total neurons (in this case 87). We then calculated the histogram of the in silico experiment using the same bins as in the empirical biological distribution. We computed a similarity score (1 – *KS_c_^M–E^*) where *KS_c_^M–E^* is the ceiled Kolmogorov-Smirnov (KS) distance between the empirical model distribution and the empirical biological distribution. The ceiled KS distance is calculated as the ratio between the actual KS distance and the maximum possible KS distance given the empirical biological distribution. We normalized the similarity score by an estimate of the similarity between empirical biological distributions (1 – *KS_c_^E–E^*) calculated by bootstrapping the empirical biological distribution. Thus, the final normalized distribution similarity score is calculated as (1 – *KS_c_^M–E^*)/(1 – *KS_c_^E–E^*). This procedure was repeated 1,000 times, each time with a different sample of artificial neurons, to estimate the uncertainty with respect to the V1 model neuronal sampling. Values reported are mean and SD.

### Statistics

Statistical significance was defined with p-value < 0.05 and the exact p-values are reported. Exact values of n and what n represents are described in the Results section and Figure captions. Reported correlations correspond to Pearson correlation coefficients. T-test was performed two-sided for two independent samples. Statistical analyses were done using the Scipy library for Python (Virtanen et al., 2020).

The ANOVA for determining the variance in the similarity scores explained by the model parameters (Supplementary Figure 4) was done using the statsmodels library for Python (Seabold & Perktold, 2010). First, for each model parameter, scores were divided in roughly equally populated bins (model depth, 4 bins; FoV, 4 values; layer depth, 6 bins; theoretical receptive field, 6 bins; number of neurons, 4 bins). The exception to this was the layer type, for which there were considerably more models with convolutional layers than pooling layers (convolutional, 632 models; pooling, 104 models). For each score type (V1 response property score or neural predictivity score), we sequentially searched for the model property that best explained the variance in the scores using a linear model without interaction terms. We started with a single model property and continued adding sequentially the remaining model parameters according to the increase in variance explained.

Data in the text is mean ± SD.

### V1 Neural Predictivity

The neuronal dataset used for the V1 neural predictivity contains extracellular recordings from 102 single-units while presenting naturalistic textures and noise images and was originally published in a study analyzing texture modulation in macaque V1 and V2 (Freeman et al., 2013). Here, we considered the stimulus-evoked responses to a subset of 315 images (20 repetitions averaged for 150ms) spanning 4deg of the visual space which constitutes the Brain-Score private dataset for the V1 neural predictivity benchmark. We considered four metrics of neural predictivity: explained variance with partial least squares (PLS) regression (25 components), explained variance with single neuron mapping, representational dissimilarity matrix (RDM), and centered kernel alignment (CKA).

For the explained variance with PLS regression we followed the same procedure as in Schrimpf et al., 2018. First, we presented the stimuli to the model after properly resizing it taking into account the model’s FoV and recorded the responses in the corresponding V1 layer. Then, we mapped the model neurons to the recorded biological neurons using a linear transformation:

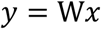

where y are the biological neuronal responses, x are model neuronal responses and W are the linear regression weights. W was calculated with a PLS regression with 25 components using the Python library scikit-learn (Pedregosa et al., 2011). The mapping procedure was implemented using a 10-fold cross-validation strategy. In each crossvalidation split, the weights are fit using training images and then used to predict the responses y’ for the held-out images. To obtain a neural predictivity score, for each biological recorded neuron, we calculated the Pearson correlation coefficient of the predicted responses y’ and the measured responses y. We then calculated the median over all the neurons normalized by their internal consistency. We used the split-half reliability as a measure of internal consistency between neural responses: we split neural responses in half across repetitions and computed the Pearson correlation coefficient between the two splits using the Spearman-Brown correction. To obtain the explained variance values reported in the Results section, we squared the normalized the neural predictivity divided by the internal consistency. Mean and standard deviations were calculated across the ten cross-validation splits.

For the explained variance using single neuron mapping, the procedure was identical with the exception that instead of using PLS regression to find the mapping between the biological neurons and the model neurons, for each single recorded neuron, we found the single neuron in the model that showed the most correlated responses.

We compared population representations using representational dissimilarity matrices (RDM) as in Kriegeskorte et al., 2008. For both the model’s activations and the V1 neuronal responses, we computed RDM according to:

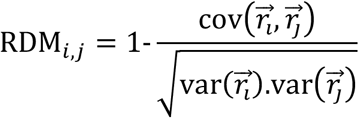

where 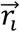 and 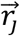 are the response vectors to the presentation of two images. We evaluated the similarity between the representations by calculating the Spearman rank correlations between the elements in the upper triangular regions of each of the RDMs. We normalized the score by an estimate of the noise ceiling, which was calculated by computing the RDM similarity between two splits of the neuronal data and corrected with the Spearman-Brown formula.

In addition to RDM, we also used linear centered kernel alignment (CKA), as a representational similarity metric. We implemented CKA according to Kornblith et al 2019:

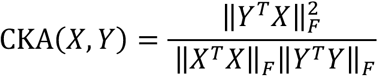

where 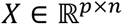 is the matrix of V1 neuronal responses of *n* neurons to *p* images and 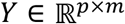 is the matrix of V1 model activations of *m* model neurons to the same *p* images. Similarly to the RDM metric, we normalized the CKA score by an estimate of the noise ceiling using split-half reliability.

### Object Recognition Behavioral Consistency

The behavioral data used in this study, as well as the corresponding analyses and comparisons with models, has been described in detail in Rajalingham et al 2015, 2018 and Schrimpf et al 2018. Here, we considered behavioral choices of human subjects performing a core object recognition behavioral paradigm. In total we used data belonging to 1,472 subjects acquired on Amazon Mechanical Turk. At each trial, an image spanning approximately 8deg was presented for 100ms followed by two response choices: one corresponding to the target object present in the image and the other, a distractor, being one of the remaining object categories. Participants responded by choosing which object was present in the image. Images consisted of objects belonging to 24 categories (10 images per category, 240 in total) displayed at different positions, orientations, and sizes, and overlayed on a random natural background. Over 300.000 responses for each target-distractor pair were obtained from multiple participants, resulting in a 240 x 24 (images x objects) response matrix when averaged across participants.

We compared the consistency between models and human behavior at four different levels: object difficulties (O1), object confusions (O2), normalized image difficulties (I1n) and normalized image confusions (I2n). For all these metrics, the procedure was analogous. First, we trained a behavioral decoder using a 24-way logistic regression (24 is the number of classes) to classify the images using the activations of the model’s penultimate layer (last layer before the 1000-way class probability layer trained on ImageNet). The behavioral decoder was trained on a separate set of 2160 images of the same object categories. We then used this regression to estimate probabilities for the test images (the ones with the behavioral data). We computed all normalized target-distractor pair probabilities, obtaining a 240 x 24 matrix (images x objects). Each entry *ij* of this matrix is given by:

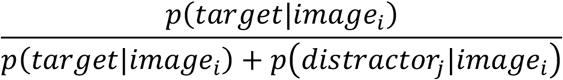

where i is the image index and j the distractor class index (different than the target index).

Prior to comparing human and model behavioral patterns, we first transform the accuracies to a discriminability measure, d’:

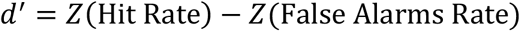

where Z is the estimated z-score of responses, Hit Rate is the accuracy of a given target-distractor pair and False Alarms Rate corresponds to how often the observers incorrectly reported seeing that target object in images where another object was present instead.

The calculation of d’ depends on the level of comparison. For O1, we calculated the discriminability of one-versus-all object choices, resulting in a vector with 24 independent values. For O2, we calculated the discriminability between each pair of object-level target-distractor, resulting in a 24×24 symmetric matrix, which we considered the off-diagonal elements on one half. For I1n, we calculated the discriminability of each image from all other objects, resulting in a vector with 240 elements. Finally, for the I2n, we calculated the discriminability of each image from each distractor object, resulting in a 5520 independent values (240 x 23). For the image-level comparisons (I1n and I2n), the response matrices were normalized by subtracting the mean Hit Rate at the corresponding level. Such normalization exposes variance unique to each image removing object-level trends.

Consistency between human and model’s discriminability was calculated with a Pearson correlation coefficient and normalized by an estimate of the noise ceiling (see Schrimpf et al 2018 for a detailed description).

### Classical V1 models

We implemented two variants of a classical V1 model containing a Gabor filter bank (GFB). The classical V1 model variant 1 consisted of an update to the V1 model frontend added to ANNs in Dapello, Marques et al., 2020. The original model contained a GFB constrained by empirical biological distributions, simple- and complex-cell nonlinearities, and a neuronal stochasticity generator. Here, we removed the neuronal stochasticity generator and added a divisive normalization stage.

We used a resolution of 224×224px and a FoV of 8deg. We implemented the GFB as a convolutional layer with a stride of four, originating a 56×56 spatial map of responses with a certain number of different channels (or cell types). Each channel in the GFB convolves the input image with a specific Gabor, parameterized by the following function:

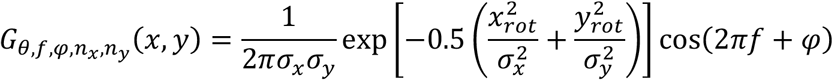

where

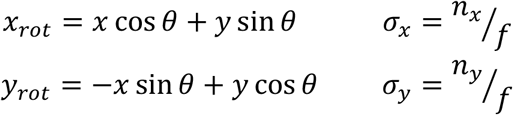

*x_rot_* and *y_rot_* are the orthogonal and parallel orientations relative to the grating edge, *θ* is the grating angular orientation, *f* is the spatial frequency of the grating, *φ* is the phase of the grating relative to the Gaussian envelope, and *σ_x_* and *σ_y_* are the standard deviations of the Gaussian envelope orthogonal and parallel to the grating edge, which can be defined as multiples (*n_x_* and *n_y_*) of the grating cycle (inverse of the frequency).

Except for the phase, *φ*, which was uniformly sampled, we relied on published V1 neurophysiology studies to generate the Gabor parameters for all of the channels. To cover this multi-dimensional space of Gabors, we considered 128 simple-cell channels and 128 complex-cell channels in the model. We used the distribution of preferred orientation in De Valois et al 1982a to sample *θ* for all the channels in the model. We used the peak spatial frequency distributions of simple- and complex-cells in De Valois et al 1982b to sample *f* for the simple- and complex-cell channels separately. We used the joint distribution of (*n_x_, n_y_*) in Ringach et al 2002 to sample these parameters for all the channels. We introduced a model hyperparameter that controls the covariance between *n_x_* and *f* (*ρ_f,n_x__*). When this value is 0, *n_x_* and *f* are sampled independently. When this value is positive, the random variables used to sample the *n_x_* and *f* distributions are positively correlated (Gabors with higher spatial frequencies will also have higher number of cycles). While the Gabors vary considerably in size, we restricted the kernel size of the Gabors to be 25×25, which corresponds to a square with 0.9deg.

After the GFB we applied two different nonlinear functions to simple- and complex-cell channels: a rectified linear transformation for simple-cells, and the spectral power of a quadrature phase-pair for complex-cells:

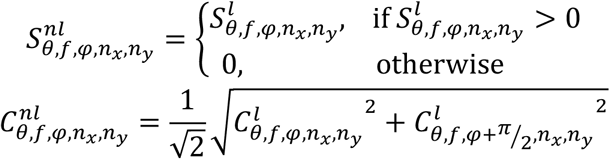

where 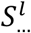 and 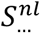 are the linear and nonlinear responses of a simple-cell channel and 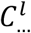 and 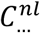 are the same for a complex-cell channel.

The final component of the V1 model variant 1 is a simple local divisive normalization stage:

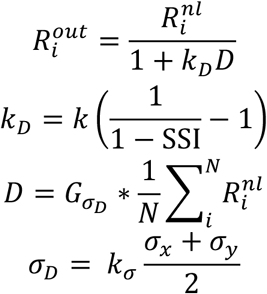

where 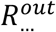 is the output response of the channel, 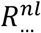 is the response after the nonlinear stage, *k_D_* is the factor controlling the magnitude of the divisive normalization and *D* is the local response activity. This local response activity is the average response across all channels convolved with a Gaussian kernel of size *σ_D_*. *k_D_* and *σ_D_* are defined separately per channel. *k_D_* is related to the surround suppression index (SSI) of the channel: for each channel we sampled the SSI distribution from Cavanaugh et al 2002 and introduced a multiplicative factor *k* common to all the channels as a model hyperparameter. We defined each channel’s *σ_D_* as the product of a hyperparameter *k_σ_* and the Gabor size (*σ_x_* and *σ_y_*).

In total, we generated 80 instantiations of the V1 model variant 1 by assigning multiple values to the three hyperparameters of the model family: *ρ_f,n_x__*, *k* and *k_σ_*. We calculated the V1 single neuron property scores and the V1 neural predictivity scores for all these models, showing the best model in Figure 7. The total number of classical V1 variant 1 models is considerably smaller than the number of ANN models considered in the same analysis (80 classical V1 variant 1 models vs 736 ANN-based V1 models).

The classical V1 model variant 2 is an implementation of the classical model used as a control in Cadena et al 2019, containing a GFB with fixed parameters and simple- and complex-cell nonlinearities. The model was implemented similarly to the classical V1 model variant 1 but the Gabor parameters were not constrained by neuronal data and, instead, were fixed to cover a range of values. The model contained 72 combinations of orientation (θ, eight different values spaced by 45deg), size (*n_x_* = *n_y_*, three values), and spatial-frequency (*f*, three values per size). We considered two phases in quadrature 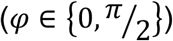 for a total of 144 simple-cell channels and the corresponding 72 complex-cell channels.

### Code availability

Code for calculating the 22 single neuron response properties, evaluating ANNs on the V1 response property similarity scores, V1 explain variance, and behavioral consistency is available in the author’s fork of the Brain-Score repository: https://github.com/tiagogmarques/brain-score

Code for creating candidate end-to-end models of the primate ventral stream from base ANNs will be available in a repository in the author’s github. Code for the classical V1 models will also be available in a dedicated repository in the author’s github.

## Supplementary Material

**Supplementary Table 1.** Empirical response properties details. Details for the 22 single neuron response properties considered in this study. Empirical distributions were obtained from the references in the last column. Histograms for all the response properties are shown in Supplementary Figure 2. For some properties there are two values in the number of neurons column. The values on top represent the total number of neurons sampled to evaluate on that property, and the numbers below in parentheses represent the number of neurons that were actually included in the neurobiological empirical distribution due to the neurons having to pass a certain criterion (see methods).

| Property | # neurons | # bins | Scale | Response Type | Reference |
| --- | --- | --- | --- | --- | --- |
| Preferred orientation | 385 | 4 | Linear | DC | DeValois et al. 1982a |
| Circular variance | 308 | 13 | Linear | DC | Ringach et al. 2002 |
| Orientation selective | 308 | 2 | Binary | DC | Ringach et al. 2002 |
| Orientation half-bandwidth | 308 | 9 | Linear | DC | Ringach et al. 2002 |
| Orthogonal / Preferred ratio | 308 | 13 | Linear | DC | Ringach et al. 2002 |
| CV / Half-bandwidth | 308 | 15 | Log. | DC | Ringach et al. 2002 |
| (Orth./Pref.) - CV | 308 | 19 | Linear | DC | Ringach et al. 2002 |
| Peak Spatial Frequency | 363 | 12 | Log. | AC or DC (baseline subtracted) | DeValois et al. 1982b |
| SF Selective | 87 | 2 | Binary | DC | Schiller et al. 1976 |
| SF Bandwidth | 87<br>(73) | 10 | Linear | DC | Schiller et al. 1976 |
| Grating Summation Field | 190<br>(148) | 8 | Log. | AC or DC (baseline subtracted) | Cavanaugh et al. 2002 |
| Surround Diameter | 190<br>(148) | 8 | Log. | AC or DC (baseline subtracted) | Cavanaugh et al. 2002 |
| Surround Suppression Index | 190 | 10 | Linear | AC or DC (baseline subtracted) | Cavanaugh et al. 2002 |
| Texture Modulation Index | 102 | 24 | Linear | Textures and Noise stimuli | Freeman, Ziemba et al. 2013 |
| Absolute Texture Modulation Index | 102 | 12 | Linear | Textures and Noise stimuli | Freeman, Ziemba et al. 2013 |
| F1/F0 ratio | 308 | 10 | Linear | AC and DC | Ringach et al. 2002 |
| Texture Selectivity | 102 | 10 | Linear | Textures stimuli | Ziemba, Freeman et al. 2016 |
| Texture Sparseness | 102 | 10 | Linear | Textures stimuli | Ziemba, Freeman et al. 2016 |
| Texture Variance Ratio | 102 | 12 | Log. | Textures stimuli | Ziemba, Freeman et al. 2016 |
| Max DC Response | 308 | 9 | Log. | DC | Ringach et al. 2002 |
| Max Texture Response | 102 | 12 | Log. | Textures stimuli | Freeman, Ziemba et al. 2013 |
| Max Noise Response | 102 | 12 | Log. | Noise stimuli | Freeman, Ziemba et al. 2013 |

**Supplementary Figure 1.**
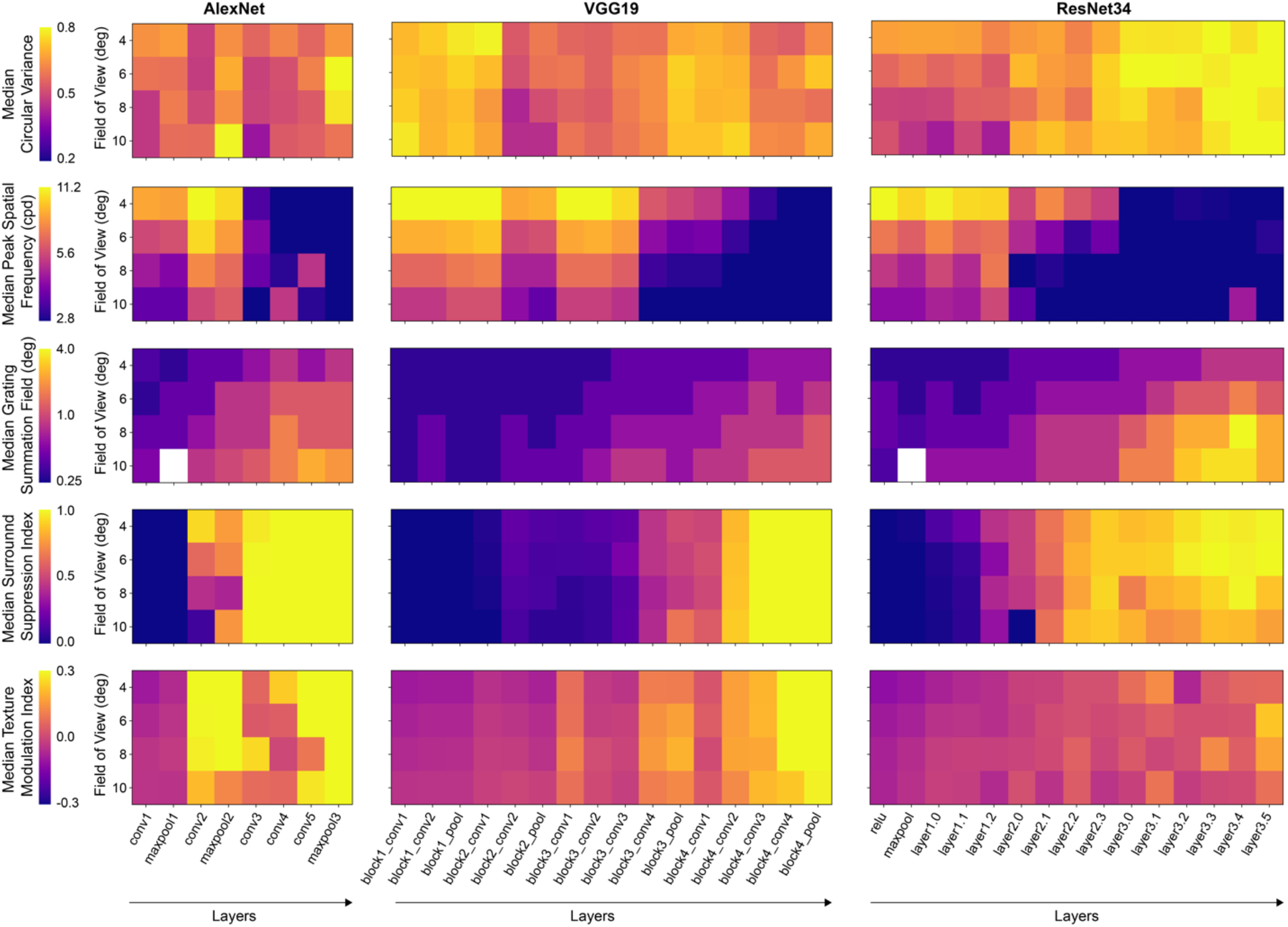
Response properties of V1 models vary with base model, field-of-view, and layer. Medians of four example response properties for the V1 models resulting from three base hierarchical models (AlexNet, VGG-19, and ResNet34). From top to bottom, circular variance, peak spatial frequency, grating summation field, surround suppression index, and texture modulation index. These examples illustrate how different response properties vary with the three steps of the model commitment: choice of base model, choice of FoV, and choice of V1-layer.

**Supplementary Figure 2.**
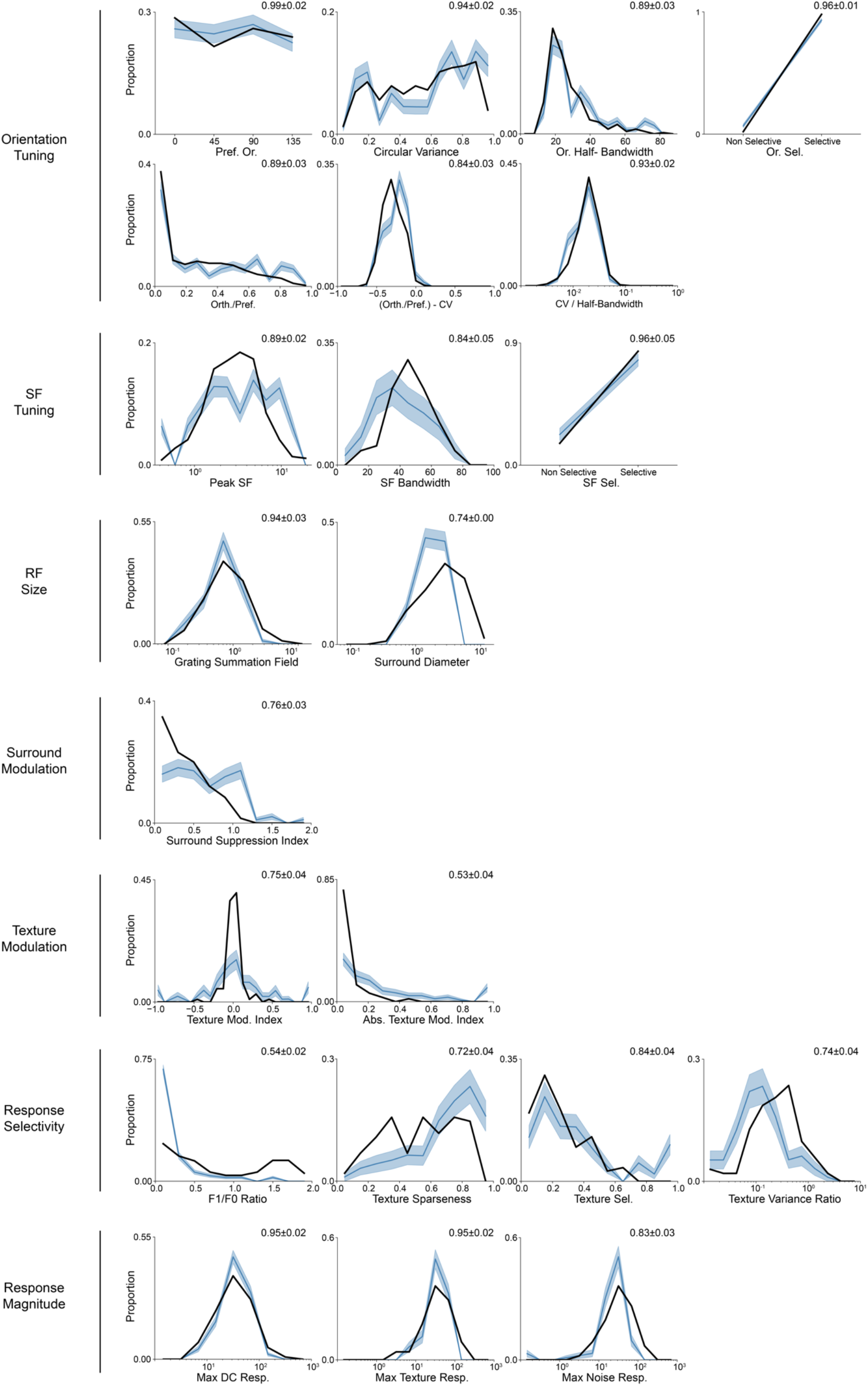
Comparison of distributions of response properties between example V1 model and macaque V1. Distributions for all the 22 single neuron response properties in macaque V1 (from published studies, black line) and an example V1 model (ResNet34-FoV8-layer2.1; obtained by performing in silico experiments in the model, thick blue line is the mean over 1,000 experiments and the shaded region is the SD). Normalized similarity score, shown in each plot at the top right corner. Response properties are organized in seven groups: orientation tuning, spatial frequency tuning, receptive field size, surround modulation, texture modulation, response selectivity, and response magnitude.

**Supplementary Figure 3.**
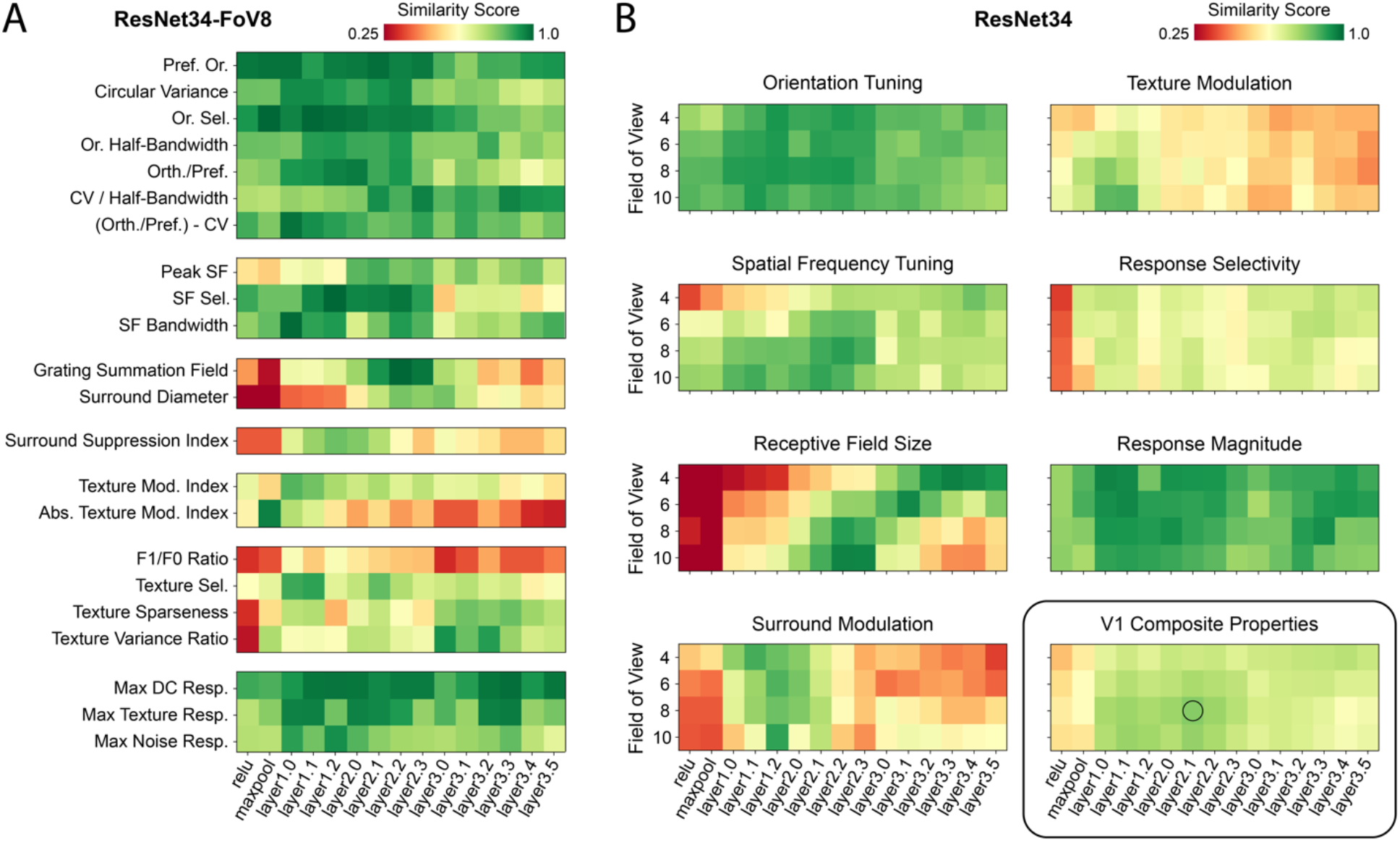
Response property similarity scores for V1 models based on the ResNet34 base model. **A.** Similarity scores for the 22 V1 response properties for all the layers of the ResNet34 with a FoV of 8 degrees. No layer is able to simultaneously approximate the distributions of all the response properties in macaque V1. Different response properties have different optimal layers. For example, grating summation field similarity score peaks at layer 2.2, while for texture modulation the score peaks at earlier layers (layer 1.0). **B.** Average scores for the seven response property groups as well as the overall V1 composite properties scores for all the V1 models based on the ResNet34 base model (all layers and FoV). For some property groups scores, there is a strong interaction between the layer and FoV. For example, for the receptive field size scores, the best models are either intermediate layers with large input FoV or higher layers with small input FoV. Best overall V1 model based on the ResNet34 base model is the one with a FoV of 8 degrees and layer 2.1 (model represented with a circle in the V1 composite properties scores and is the same model as in Figure 2, and 3).

**Supplementary Figure 4.**
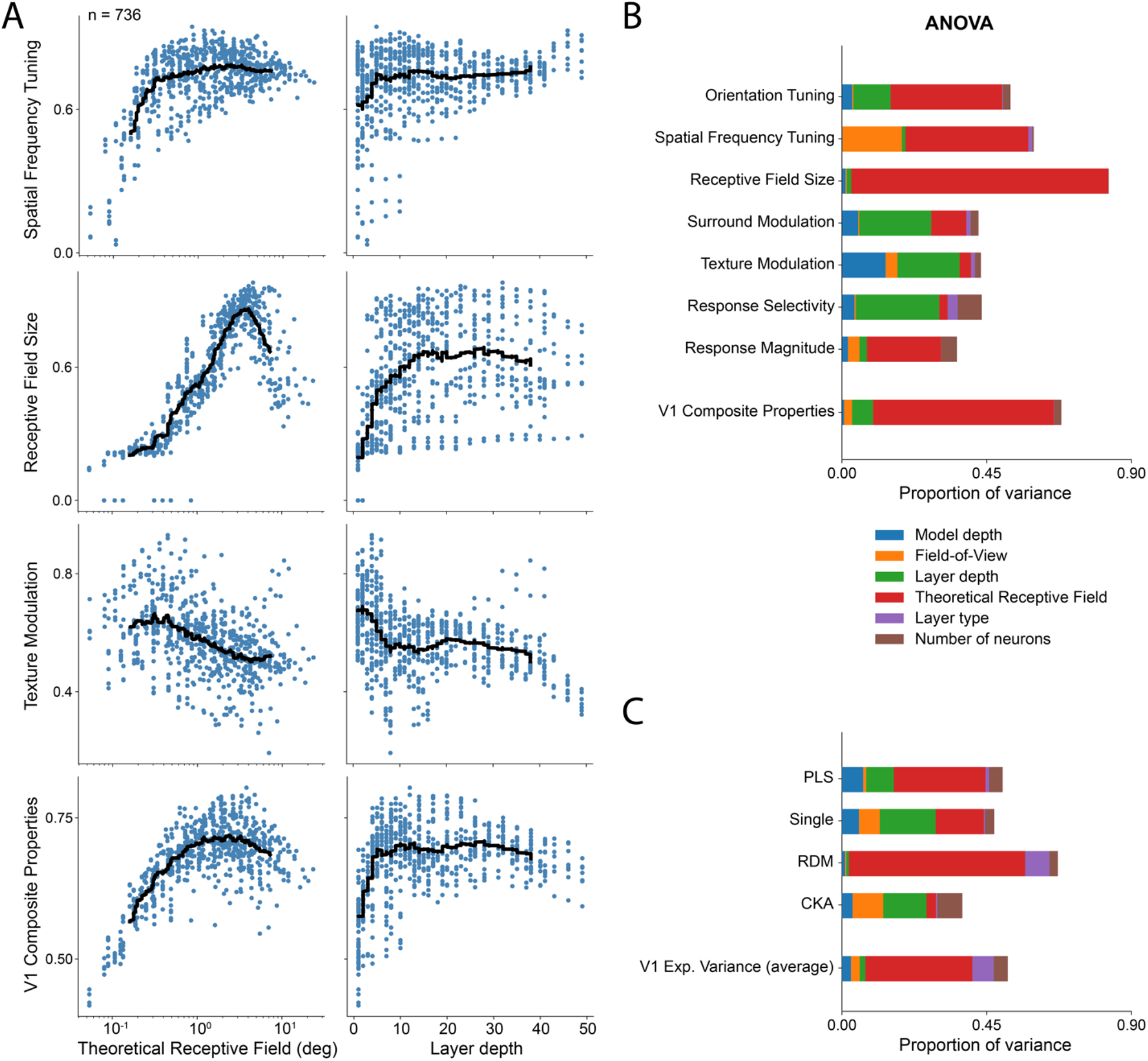
Model properties account for the majority of the variance in the response property scores. **A.** Similarity scores for three example response property groups (spatial frequency tuning, receptive field size, and texture modulation) and V1 composite properties as a function of the model’s theoretical receptive field size and layer depth for all the V1 ANN-based models (n=736). Black line is the moving average (window of 100). Depending on the response properties, over all the models, V1 similarity scores vary with model properties. For example, receptive field size similarity scores are optimal for models with theoretical receptive field sizes of ~4 degrees. **B.** Sequential ANOVA for determining how much variance in the property scores can be explained by model properties excluding interactions (see methods). For the V1 composite properties scores, most of the variance (67.3%) can be explained by six model properties: model depth, FoV, layer depth, theoretical receptive field, layer type and number of neurons. The theoretical receptive field corresponds to the fraction of the visual input that neurons can be influenced by (in degrees) and does not correspond to the receptive field size as determined by the *in silico* electrophysiology experiments. Layer depth is defined as the cumulative number of non-linear processing stages until the model layer and model depth is the total number of non-linear processing stages in the ANN. Layer type can be either convolution followed by a ReLU or a maxpool operation. Theoretical receptive field size is the model property that explains most variance in the V1 composite and property groups scores, followed by the layer depth. **C.** Same as B but for different V1 neural predictivity metrics in a given neuronal dataset.

**Supplementary Figure 5.**
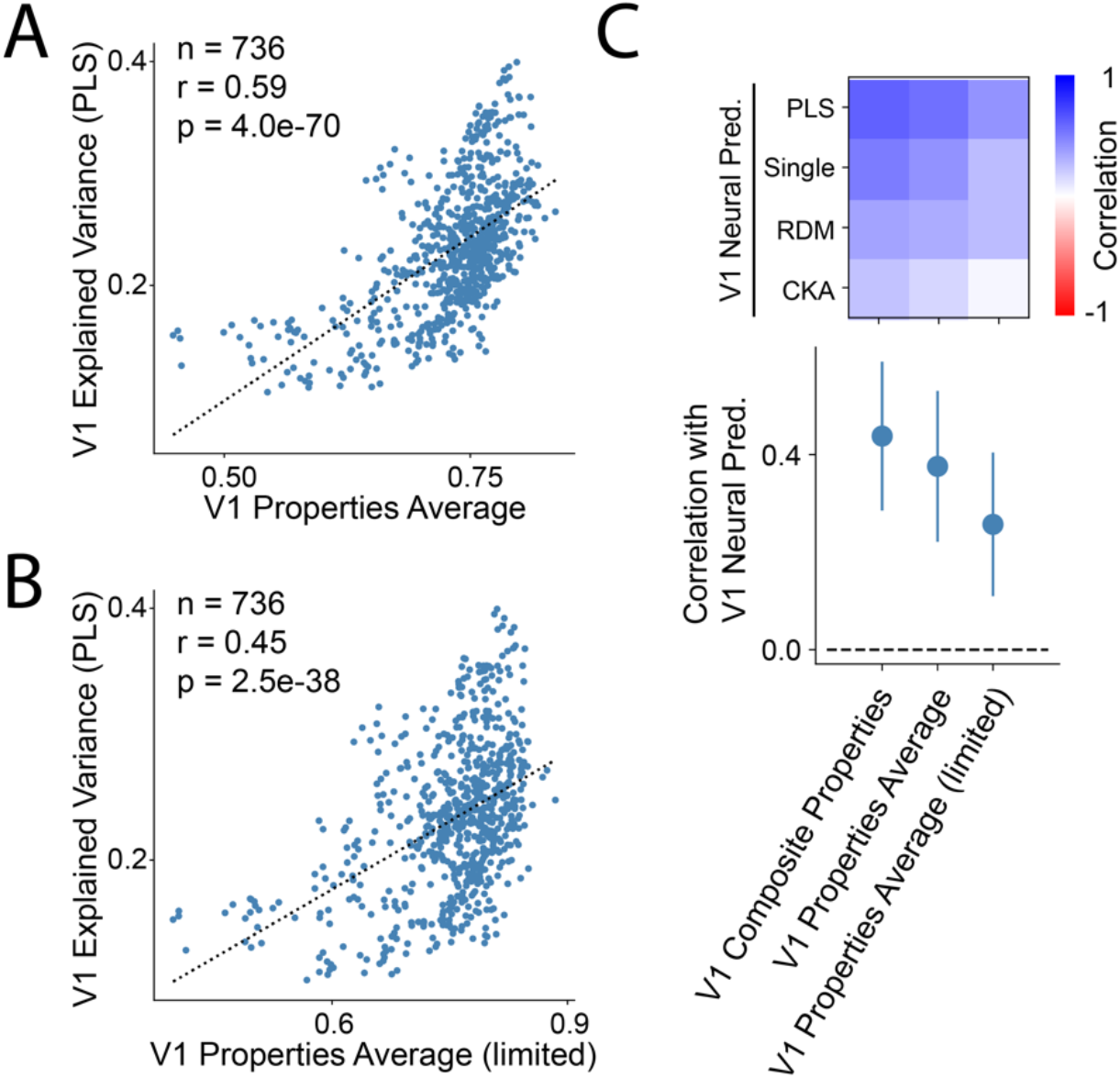
Correlation of model’s property distribution similarity scores with neural predictivity is robust. **A.** Similar to Figure 5 A but V1 property scores are averaged directly along the 22 response properties and not pre-pooled in groups. Comparison of model’s ability to explain variance in macaque V1 responses in a neuronal dataset (Freeman, Ziemba et al 2013) using PLS regression and V1 properties average scores (across 736 ANN V1 models). Model’s explained variance is positively correlated with the V1 properties average scores. **B.** Same as **A** but discarding the response property scores that are calculated using the same stimuli and neuronal responses that are used for the V1 explained variance: texture modulation index, absolute texture modulation index, texture selectivity, texture sparseness, variance ratio, maximum texture response, and maximum noise response. There is still a significant correlation between the V1 explained variance and the average of the limited pool of non-overlapping response property scores (15 response properties). **C.** Top, correlation of the three distinct ways to pool the V1 property scores (composite, average, and limited average with different V1 neural predictivity metrics (explained variance with PLS regression and single neuron mapping, and representational similarity analysis with RDM and CKA). Bottom, same as above but showing the mean and SD across the different neural predictivity metrics.

## Notes

### Competing Interest Statement

The authors have declared no competing interest.

## References

Adelson, E. H., & Bergen, J. R. (1985). Spatiotemporal energy models for the perception of motion. Journal of the Optical Society of America A, Optics and Image Science, 2(2), 284–299. https://doi.org/10.1364/JOSAA.2.000284

Arend, L., Han, Y., Schrimpf, M., Bashivan, P., Kar, K., Poggio, T., … Boix, X. (2018). Single units in a deep neural network functionally correspond with neurons in the brain: preliminary results. CBMM Memo, (093), 1–23.

Bair, W., Cavanaugh, J. R., & Movshon, J. A. (2003). Time course and time-distance relationships for surround suppression in macaque V1 neurons. Journal of Neuroscience, 23(20), 7690–7701. https://doi.org/23/20/7690 [pii]

Bashivan, P., Kar, K., & DiCarlo, J. J. (2019). Neural population control via deep image synthesis. In Science (Vol. 364). https://doi.org/10.1126/science.aav9436

Brendel, W., & Bethge, M. (2019). Approximating CNNs with Bag-of-local-Features models works surprisingly well on ImageNet, 1–15.

Cadena, S. A., Denfield, G. H., Walker, E. Y., Gatys, L. A., Tolias, A. S., Bethge, M., & Ecker, A. S. (2019). Deep convolutional models improve predictions of macaque V1 responses to natural images Author summary. PLoS Computational Biology, 1–28. https://doi.org/10.12751/g-node.2e31e3

Cadieu, C. F., Hong, H., Yamins, D. L. K., Pinto, N., Ardila, D., Solomon, E. A., … DiCarlo, J. J. (2014). Deep Neural Networks Rival the Representation of Primate IT Cortex for Core Visual Object Recognition. PLoS Computational Biology, 10(12). https://doi.org/10.1371/journal.pcbi.1003963

Carandini, M., Heeger, D. J., & Movshon, J. A. (1997). Linearity and normalization in simple cells of the macaque primary visual cortex. The Journal of Neuroscience : The Official Journal of the Society for Neuroscience, 17(21), 8621–8644.

Cavanaugh, J. R., Bair, W., & Movshon, J. A. (2002). Nature and Interaction of Signals From the Receptive Field Center and Surround in Macaque V1 Neurons. Journal of Neurophysiology, 88(5), 2530–2546. https://doi.org/10.1152/jn.00692.2001

Chollet, F. (2015). Keras.

Dapello, J., Marques, T., Schrimpf, M., Geiger, F., Cox, D. D., & DiCarlo, J. J. (2020). Simulating a primary visual cortex at the front of CNNs improves robustness to image perturbations. NeurIPS, 1–30. https://doi.org/10.1101/2020.06.16.154542

De Valois, R. L., Albrecht, D. G., & Thorell, L. G. (1982). Spatial Frequency Selectivity of Cells in Macaque Visual Cortex. Vision Research, 22, 545–559.

De Valois, R. L., Yund, E. W., & Hepler, N. (1982). The orientation and direction selectivity of cells in macaque visual cortex. Vision Research, 22, 531–544.

DiCarlo, J. J., & Cox, D. (2007). Untangling invariant object recognition. Trends in Cognitive Sciences, 11(8), 333–341. https://doi.org/10.1016/j.tics.2007.06.010

DiCarlo, J. J., Zoccolan, D., & Rust, N. C. (2012). How Does the Brain Solve Visual Object Recognition? Neuron, 73(3), 415–434. https://doi.org/10.1016/j.neuron.2012.01.010

Engstrom, L., Ilyas, A., Santurkar, S., & Tsipras, D. (2019). Robustness (Python Library).

Fabre-Thorpe, M., Richard, G., & Thorpe, S. J. (1998). Rapid categorization of natural images by rhesus monkeys. NeuroReport, 9(2), 303–308. https://doi.org/10.1097/00001756-199801260-00023

Felleman, D. J., & Van Essen, D. C. (1991). Distributed hierarchical processing in the primate cerebral cortex. Cerebral Cortex, 1(1), 1–47.

Freeman, J., Ziemba, C. M., Heeger, D. J., Simoncelli, E. P., & Movshon, J. A. (2013). A functional and perceptual signature of the second visual area in primates. Nature Neuroscience, 16(7), 974–981. https://doi.org/10.1038/nn.3402

Geirhos, R., Rubisch, P., Michaelis, C., Bethge, M., Wichmann, F. A., & Brendel, W. (2019). ImageNet-trained CNNs are biased towards texture; increasing shape bias improves accuracy and robustness. In ICLR (pp. 1–22).

Güçlü, U., & van Gerven, M. A. J. (2015). Deep Neural Networks Reveal a Gradient in the Complexity of Neural Representations across the Brain’s Ventral Visual Pathway. The Journal of Neuroscience, 35(27), 10005–10014. https://doi.org/10.1523/JNEUROSCI.5023-14.2015

He, K., Zhang, X., Ren, S., & Sun, J. (2016). Deep Residual Learning for Image Recognition. In CVPR (pp. 1–12).

Heeger, D. J., Simoncelli, E. P., & Movshon, J. A. (1996). Computational models of cortical visual processing. Proceedings of the National Academy of Sciences of the United States of America, 93(2), 623–627. https://doi.org/10.1073/pnas.93.2.623

Helland, I. (2006). Partial Least Squares Regression. In Encyclopedia of Statistical Sciences. American Cancer Society. https://doi.org/10.1002/0471667196.ess6004.pub2

Horwitz, G. D., & Hass, C. A. (2012). Nonlinear analysis of macaque V1 color tuning reveals cardinal directions for cortical color processing. Nature Neuroscience, 15(6), 913–919. https://doi.org/10.1038/nn.3105

Jones, H. E., Grieve, K. L., Wang, W., & Sillito, A. M. (2001). Surround suppression in primate V1. Journal of Neurophysiology, 86(4), 2011–2028. https://doi.org/10.1152/jn.2001.86.4.2011

Jones, J. P., & Palmer, L. A. (1987). The two-dimensional spatial structure of simple receptive fields in cat striate cortex. Journal of Neurophysiology, 58(6), 1187–1211. https://doi.org/10.1152/jn.1987.58.6.1187

Kapadia, M. K., Ito, M., Gilbert, C. D., & Westheimer, G. (1995). Improvement in visual sensitivity by changes in local context: Parallel studies in human observers and in V1 of alert monkeys. Neuron, 15(4), 843–856. https://doi.org/10.1016/0896-6273(95)90175-2

Kar, K., Kubilius, J., Schmidt, K., Issa, E. B., & DiCarlo, J. J. (2019). Evidence that recurrent circuits are critical to the ventral stream’s execution of core object recognition behavior. Nature Neuroscience. https://doi.org/10.1038/s41593-019-0392-5

Khaligh-Razavi, S. M., & Kriegeskorte, N. (2014). Deep Supervised, but Not Unsupervised, Models May Explain IT Cortical Representation. PLoS Computational Biology, 10(11). https://doi.org/10.1371/journal.pcbi.1003915

Kietzmann, T. C., Spoerer, C. J., Sörensen, L. K. A., Cichy, R. M., & Hauk, O. (2019). Recurrence is required to capture the representational dynamics of the human visual system, 116(43). https://doi.org/10.1073/pnas.1905544116

Kornblith, S., Norouzi, M., Lee, H., & Hinton, G. (2019). Similarity of Neural Network Representations Revisited.

Kriegeskorte, N. (2015). Deep Neural Networks: A New Framework for Modeling Biological Vision and Brain Information Processing. Annual Review of Vision Science, 1(1), 417–446. https://doi.org/10.1146/annurev-vision-082114-035447

Kriegeskorte, N., Mur, M., Ruff, D. A., Kiani, R., Bodurka, J., Esteky, H., … Bandettini, P. A. (2008). Matching Categorical Object Representations in Inferior Temporal Cortex of Man and Monkey. Neuron, 60(6), 1126–1141. https://doi.org/10.1016/j.neuron.2008.10.043

Krizhevsky, A., Sutskever, I., & Geoffrey E., H. (2012). ImageNet Classification with Deep Convolutional Neural Networks. In NIPS (pp. 1097–1105). https://doi.org/10.1109/5.726791

Kubilius, J., Schrimpf, M., Kar, K., Hong, H., Majaj, N. J., Rajalingham, R., … DiCarlo, J. J. (2019). Brain-Like Object Recognition with High-Performing Shallow Recurrent ANNs. NeurIPS, (NeurIPS), 1–12.

Kubilius, J., Schrimpf, M., Nayebi, A., Bear, D., Yamins, D. L. K., & Dicarlo, J. J. (2018). CORnet : Modeling the Neural Mechanisms of Core Object Recognition. BioRxiv, 1–9. https://doi.org/10.1101/408385

Lamme, V. A. F. (1995). The Neurophysiology of Figure-Ground Segregation in Primary Visual Cortex. The Journal of Neuroscience, 15(2), 1605–1615.

Laskar, M. N. U., Giraldo, L. G. S., & Schwartz, O. (2018). Correspondence of Deep Neural Networks and the Brain for Visual Textures. ArXiv, 1–17.

Madry, A., Makelov, A., Schmidt, L., Tsipras, D., & Vladu, A. (2019). Towards Deep Learning Models Resistant to Adversarial Attacks. ArXiv, 1–28.

Matteucci, G., Marotti, R. B., Riggi, M., Rosselli, F. B., & Zoccolan, D. (2019). Nonlinear processing of shape information in rat lateral extrastriate cortex. Journal of Neuroscience, 39(9), 1649–1670. https://doi.org/10.1523/JNEUROSCI.1938-18.2018

Mishkin, M., Ungerleider, L. G., & Macko, K. A. (1983). Object vision and spatial vision: two cortical pathways. Trends in Neurosciences, 6(C), 414–417. https://doi.org/10.1016/0166-2236(83)90190-X

Nassi, J. J., Lomber, S. G., & Born, R. T. (2013). Corticocortical Feedback Contributes to Surround Suppression in V1 of the Alert Primate. Journal of Neuroscience, 33(19), 8504–8517. https://doi.org/10.1523/JNEUROSCI.5124-12.2013

Nayebi, A., Bear, D., Kubilius, J., Kar, K., Ganguli, S., Sussillo, D., … Yamins, D. L. K. (2018). Task-Driven Convolutional Recurrent Models of the Visual System. NeurIPS. https://doi.org/arXiv:1807.00053v2

Nurminen, L., Merlin, S., Bijanzadeh, M., Federer, F., & Angelucci, A. (2018). Top-down feedback controls spatial summation and response amplitude in primate visual cortex. Nature Communications, 9(1). https://doi.org/10.1038/s41467-018-04500-5

Paszke, A., Gross, S., Massa, F., Lerer, A., Bradbury, J., Chanan, G., … Chintala, S. (2019). PyTorch: An imperative style, high-performance deep learning library. NeurIPS.

Pedregosa, F., Varoquaux, G., Gramfort, A., Michel, V., Thirion, B., Grisel, O., … Duchesnay, E. (2011). Scikit-learn: Machine Learning in Python. Journal of Machine Learning Research, 12, 2825–2830.

Rajalingham, R., Issa, E. B., Bashivan, P., Kar, K., Schmidt, K., & DiCarlo, J. J. (2018). Large-scale, high-resolution comparison of the core visual object recognition behavior of humans, monkeys, and state-of-the-art deep artificial neural networks. The Journal of Neuroscience, 38(33), 7255–7269. https://doi.org/10.1523/JNEUROSCI.0388-18.2018

Rajalingham, R., Schmidt, K., & DiCarlo, J. J. (2015). Comparison of Object Recognition Behavior in Human and Monkey. Journal of Neuroscience, 35(35), 12127–12136. https://doi.org/10.1523/JNEUROSCI.0573-15.2015

Richards, B. A., Lillicrap, T. P., Beaudoin, P., Bengio, Y., Bogacz, R., Christensen, A., … Kording, K. P. (2019). A deep learning framework for neuroscience. Nature Neuroscience, 22(11), 1761–1770. https://doi.org/10.1038/s41593-019-0520-2

Ringach, D. L., Shapley, R. M., & Hawken, M. J. (2002). Orientation selectivity in macaque V1: diversity and laminar dependence. The Journal of Neuroscience, 22(13), 5639–5651. https://doi.org/20026567

Saxe, A., Nelli, S., & Summerfield, C. (2021). If deep learning is the answer, what is the question? Nature Reviews Neuroscience, 22(1), 55–67. https://doi.org/10.1038/s41583-020-00395-8

Sceniak, M. P., Ringach, D. L., Hawken, M. J., & Shapley, R. (1999). Contrast’s effect on spatial summation by macaque V1 neurons. Nature Neuroscience, 2(8), 733–739. https://doi.org/10.1038/11197

Schiller, P. H., Finlay, B. L., & Volman, S. F. (1976). Quantitative studies of single-cell properties in monkey striate cortex. III. Spatial Frequency. Journal of Neurophysiology, 39(6), 1334–1351. https://doi.org/10.1152/jn.1976.39.6.1352

Schiller, Peter H, Finlay, L., & Volman, F. (1976). Quantitative Studies of Single-Cell Properties in Monkey Striate Cortex. II. Orientation Specificity and Ocular Dominance. Journal of Neurophysiology, 39(6), 1320–1333. https://doi.org/018.030.060.187

Schrimpf, M., Kubilius, J., Hong, H., Majaj, N. J., Rajalingham, R., Issa, E. B., … DiCarlo, J. J. (2018). Brain-Score: Which Artificial Neural Network for Object Recognition is most Brain-Like? BioRxiv, 1–9. https://doi.org/10.1101/407007

Schrimpf, M., Kubilius, J., Lee, M. J., Ratan Murty, N. A., Ajemian, R., & DiCarlo, J. J. (2020). Integrative Benchmarking to Advance Neurally Mechanistic Models of Human Intelligence. Neuron, 1–11. https://doi.org/10.1016/j.neuron.2020.07.040

Seabold, S., & Perktold, J. (2010). Statsmodels: Econometric and Statistical Modeling with Python. Proceedings of the 9th Python in Science Conference, (Scipy), 92–96. https://doi.org/10.25080/majora-92bf1922-011

Serre, T. (2019). Deep Learning: The Good, the Bad, and the Ugly. Annual Review of Vision Science, 5(1), 399–426. https://doi.org/10.1146/annurev-vision-091718-014951

Simonyan, K., & Zisserman, A. (2015). Very Deep Convolutional Networks for Large-Scale Image Recognition. In ICLR (pp. 1–14). https://doi.org/10.2146/ajhp170251

Skottun, B. C., De Valois, R. L., Grosof, D. H., Movshon, J. A., Albrecht, D. G., & Bonds, A. B. (1991). Classifying simple and complex cells on the basis of response modulation. Vision Research, 31(7–8), 1079–1086. https://doi.org/10.1016/0042-6989(91)90033-2

Tang, H., Schrimpf, M., Lotter, W., Moerman, C., Paredes, A., Ortega Caro, J., … Kreiman, G. (2018). Recurrent computations for visual pattern completion. Proceedings of the National Academy of Sciences, 115(35), 8835–8840. https://doi.org/10.1073/pnas.1719397115

Virtanen, P., Gommers, R., Oliphant, T. E., Haberland, M., Reddy, T., Cournapeau, D., … Vázquez-Baeza, Y. (2020). SciPy 1.0: fundamental algorithms for scientific computing in Python. Nature Methods, 17(3), 261–272. https://doi.org/10.1038/s41592-019-0686-2

Yamins, D. L. K., & DiCarlo, J. J. (2016). Using goal-driven deep learning models to understand sensory cortex. Nature Neuroscience, 19(3), 356–365. https://doi.org/10.1038/nn.4244

Yamins, D. L. K., Hong, H., Cadieu, C. F., Solomon, E. A., Seibert, D., & DiCarlo, J. J. (2014). Performance-optimized hierarchical models predict neural responses in higher visual cortex. Proceedings of the National Academy of Sciences, 111(23), 8619–8624. https://doi.org/10.1073/pnas.1403112111

Zhou, H., Friedman, H. S., von R, der H., & von der Heydt, R. (2000). Coding of border ownership in monkey visual cortex. Journal of Neuroscience, 20(17), 6594. https://doi.org/10.1523/JNEUROSCI.2797-12.2013

Ziemba, C. M., Freeman, J., Movshon, J. A., & Simoncelli, E. P. (2016). Selectivity and tolerance for visual texture in macaque V2. Proceedings of the National Academy of Sciences, 113(22), E3140–E3149. https://doi.org/10.1073/pnas.1510847113

